# Insights into the Translational Activation Mechanisms of the *COX1* mRNA in Yeast Mitochondria

**DOI:** 10.1101/2024.11.01.621605

**Authors:** Angelica Zamudio-Ochoa, Yolanda Camacho-Villasana, Aldo E. García-Guerrero, Xochitl Pérez-Martínez

## Abstract

Mitochondrial translation is a critical regulatory step in mitochondrial genome expression. In *Saccharomyces cerevisiae*, translational activators are believed to bind to the 5’ UTRs of their target mRNAs to position the mitochondrial ribosome at the start codon. Pet309 and Mss51 are translational activators of *COX1* mRNA, which encodes subunit one of cytochrome *c* oxidase. Pet309 physically interacts with *COX1* mRNA, but no direct interaction of Mss51 with its target mRNA has been detected. Currently, the mechanisms underlying translational activation of *COX1*, or any other mitochondrial gene, remain poorly understood.

To explore in depth the mechanism of *COX1* mRNA translational activation, we studied the association of Pet309 and Mss51 with the mitochondrial ribosome. Both Pet309 and Mss51 interact with the mitoribosome regardless of the presence of *COX1* mRNA or of each other. Pet309’s association with the ribosome and with *COX1* mRNA depends on its N-terminal domain. These findings indicate that Pet309 and Mss51 stably interact with the mitoribosome independently of an active translation. By integrating our data with previously published research, we propose a new mechanism of *COX1* mRNA translation activation.

**SUMMARY STATEMENT:** Yeast mitochondrial mRNAs require translational activators by an almost unknown mechanism. Based on our findings on Mss51 and Pet309 function, we present a new model for translation of the *COX1* mRNA

## INTRODUCTION

Since the discovery of the mitochondrial genome in the 1960s (Nass and Nass, 1963, Schatz et al., 1964), extensive research has been conducted to unravel the molecular mechanisms underlying its maintenance, expression, and regulation. Among these mechanisms, translation stands out as a critical regulatory step in mitochondrial genome expression (Couvillion et al., 2016, Fontanesi, 2013, Richter-Dennerlein et al., 2016). Despite significant progress, our understanding of this process remains incomplete. Most studies on mitochondrial translation use the yeast *Saccharomyces cerevisiae* as a model. In *S. cerevisiae*, the mitochondrial genome encodes 2 ribosomal RNAs, 22 transfer RNAs, the 9S RNA (a component of RNase P), and eight proteins (Foury et al., 1998). Seven of these mitochondria-encoded proteins are integral subunits of the respiratory complexes and ATP synthase, and their translation is tightly regulated and synchronized with the assembly of the nuclear-encoded subunits (Couvillion et al., 2016).

Yeast mitochondrial mRNAs feature long 5’ untranslated regions (UTRs), essential for translation regulation via translational activators, though the precise regulation mechanism remains unclear. It is hypothesized that these translational activators help position mitochondrial ribosomes on the start codon of the mRNA (Fox, 1996). Each mitochondrial mRNA has a unique set of translational activators (Herrmann et al., 2013). Furthermore, all yeast mitochondrial mRNAs depend on at least one translational activator containing pentatricopeptide repeat (PPR) motifs, which function as RNA-binding domains (Lipinski et al., 2011). PPR proteins belong to the Alpha-Solenoids structures tandem repeat proteins group (STRPs) (Arrias et al., 2024). PPR proteins consists of multiple 35-amino-acids degenerate repeats, with each repeat forming two antiparallel α-helices (Ringel et al., 2011). Each repeat binds one nucleotide of its target mRNA for specific RNA recognition (Filipovska and Rackham, 2013). Currently, 15 PPR proteins have been identified in yeast, all of which are localized within mitochondria and are essential in in RNA metabolism functions, including transcription, stability, and translation of mRNAs (Herbert et al., 2013, Lipinski et al., 2011).

The PPR protein Pet309 is a translational activator of the *COX1* mRNA (Manthey and McEwen, 1995). This mRNA encodes Cox1, the largest and core subunit of Cytochrome *c* Oxidase (C*c*O) (Scott, 1995). C*c*O is indispensable for respiration, serving as the terminal electron acceptor in the electron transport chain and catalyzing the reduction of molecular oxygen to water. Pet309 specifically acts on the 5’UTR of the *COX1* mRNA to activate its translation. We previously reported the cooperative action of Pet309’s PPR repeats on *COX1* mRNA binding (Zamudio-Ochoa et al., 2014). Furthermore, Pet309, along with other translational activators, has been identified in mitochondrial ribosome co-purifications (Kehrein et al., 2015).

In addition to Pet309, Mss51 is also a *COX1* mRNA translational activator, despite lacking a detectable physical interaction with the transcript (Zamudio-Ochoa et al., 2014). Mss51 coordinates the synthesis of Cox1 with its assembly into the C*c*O complex (Perez-Martinez et al., 2003). Mss51 physically interacts with the newly synthesized Cox1 and acts as a chaperone, facilitating its assembly into C*c*O. The interaction between Cox1 and Mss51 leads to the formation of translational repressor complexes known as COA complexes (Fontanesi et al., 2011, Mick et al., 2010, Pierrel et al., 2007). These complexes serve as checkpoints during C*c*O assembly, halting the translational activator function of Mss51. When C*c*O assembly is completed, Mss51 is released from the COA complexes, initiating a new round of *COX1* mRNA translation. However, if defects in C*c*O assembly occur, Mss51 remains sequestered in the COA complexes, preventing *COX1* mRNA translation (For review (Franco et al., 2020)). Notably, the mammalian Mss51 orthologue is a potential target for obesity and type 2 diabetes prevention (Ali et al., 2024, Moyer and Wagner, 2015).

Despite extensive efforts to elucidate the roles of Pet309 and Mss51 as translational activators, the mechanisms underlying their activity in *COX1* translation remain elusive. Furthermore, the interplay between these two activators in regulating Cox1 synthesis is poorly understood. Contrary to the proven association of Pet309 with the mitoribosome, studies on Mss51’s association are inconsistent: some studies found Mss51 associated with high molecular weight complexes (Fontanesi et al., 2010, Verma et al., 2021), while another study did not detect it in purified mitoribosomes (Kehrein et al., 2015). To advance our understanding of *COX1* mRNA translation, we investigated Pet309 and Mss51’s interactions with the mitoribosome. Using sucrose gradient ultracentrifugation, we found that Pet309 and Mss51 independently bind the mitochondrial ribosome regardless of *COX1* mRNA active translation or each other’s presence. In addition, we identified the N-terminal region of Pet309 as critical for ribosome and *COX1* mRNA binding. Based on these findings and previous data from other groups, we propose a new model for *COX1* mRNA translation.

## RESULTS

### Pet309 constitutively interacts with the mitoribosome independently of the *COX1* mRNA and Mss51

Translational activators like Pet309 are proposed to position the mitoribosome on the mRNA start codon of its target mRNA (Fox, 1996). Indeed, Pet309 has shown genetical and physical interaction with the *COX1* mRNA (Manthey and McEwen, 1995, Zamudio-Ochoa et al., 2014). In addition, Pet309 is a component of the MIOREX complex and associates with the mitoribosome (Kehrein et al., 2015). We looked closer into this interaction and investigated if the Pet309-mitoribosome association depended on the presence of the *COX1* mRNA and Mss51. To analyze the Pet309-mitoribosome interaction, Pet309 was tagged with three hemagglutinin epitopes at its C-terminal domain (*PET309-3xHA*). This construct was cloned into a centromeric plasmid (*CEN*) and expressed it episomally in a *pet309*Δ strain. No respiratory defect in the strain bearing the *PET309-3xHA* construct was observed, as previously reported (Tavares-Carreon et al., 2008, Zamudio-Ochoa et al., 2014). To evaluate Pet309 association with the mitoribosome, mitochondria isolated from the *PET309-3xHA* strain were lysed using 0.1% digitonin, and the clarified lysate was loaded onto a discontinuous sucrose gradient (Garcia-Guerrero et al., 2018). After ultracentrifugation, the gradient was divided into seven fractions and analyzed by western blot. Pet309-3xHA was detected in the same fractions as bS1 (Mrp51 – old nomenclature) and uL23 (Mrp20 – old nomenclature), mitoribosomal proteins of the small and large subunits, respectively (Figure 1A). As expected, citrate synthase (CS), an enzyme from the citric acid cycle unrelated to the respiratory chain, was found in the surface portion of the gradient. Notably, Pet309-3xHA was not detected in top fractions, suggesting that all Pet309 present in mitochondria interacts with the mitoribosome. Interestingly, Pet309-3xHA was almost undetectable by western blot in a strain lacking mitochondrial DNA (*ρ°*), in which mitochondrial DNA and, therefore, mitoribosomal RNAs are absent, suggesting that stability of Pet309 depends on the presence of mitoribosomes and/or mitochondrial DNA (Figure 1B). To have more insights on Pet309 and its interaction with the mitoribosome, we aimed to analyze Pet309-3xHA migration in a *ρ°* strain. Due to the observed instability of Pet309 in a *ρ°* strain, we cloned *PET309-3xHA* into a 2 μ plasmid for overexpression (*PET309-3xHA^OE^*). In contrast to *CEN*-plasmid expression, Pet309-3xHA^OE^ was detected on the surface fraction of the sucrose gradient (Figure 1C). This finding indicates that Pet309 migration into high molecular weight fractions depends on the presence of the mitoribosome.

**Figure 1.**
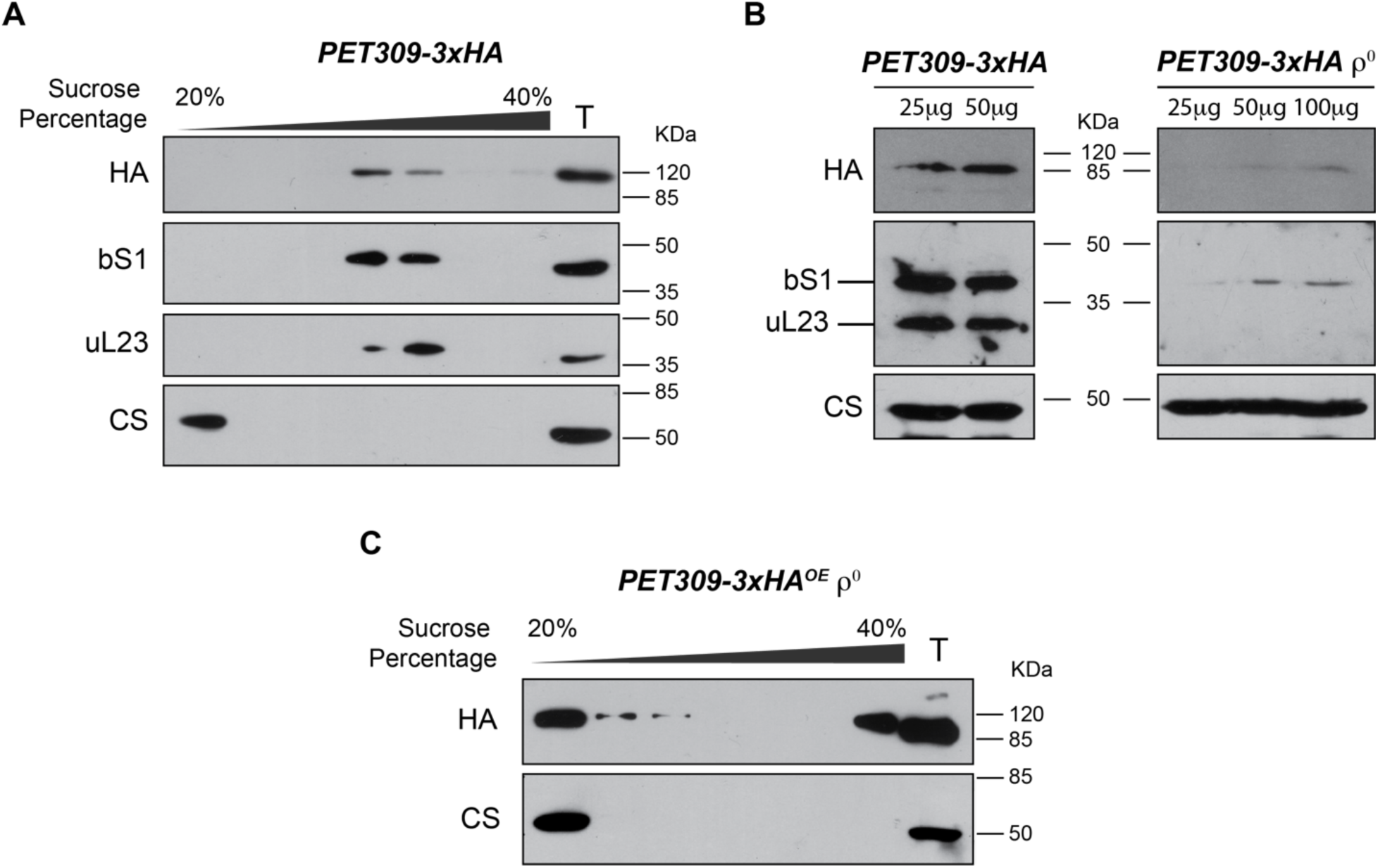
Pet309 associates to the mitoribosome. A) 500 μg of mitochondrial protein from the *PET309-3xHA* strain were lysed with 1% digitonin. Clarified lysate was centrifuged in a discontinuous 20 to 40% sucrose gradient. Seven fractions were collected and analyzed by SDS-PAGE and western blot using the indicated antibodies. B) Western blot analysis with indicated amounts of mitochondrial protein from the *PET309-3xHA* and *PET309-3xHA ρ^0^* strains. C) Same experiment shown in A) with lysed mitochondria from the *ρ^0^* strain. For efficient immunodetection, a strain overexpressing *PET309-3xHA^OE^* was used in this experiment. In all experiments citrate synthase (CS) was used as loading control or a protein independent form the respiratory chain function. bS1, small mitoribosomal subunit (Mrp51); uL23 large mitoribosomal subunit (Mrp21); T, total fraction equivalent to 7% from the loaded clarified lysate. Uncropped blots in Figure S4.

Pet309 physically interacts with the *COX1* mRNA (Zamudio-Ochoa et al., 2014), thus Pet309 interaction with the mitoribosome could be mediated by an active translation of its specific mRNA, as previously observed for the *COB* translational activator, Cbs2 (Krause-Buchholz et al., 2004). To determine if Pet309 association with the mitoribosome rely upon the presence of the *COX1* mRNA, we expressed Pet309-3xHA in a strain that lacks the entire *COX1* ORF, including its UTRs (*cox1*Δ). In this strain the *COX1* gene was deleted from 787 base pairs (bp) upstream of the *AUG* start codon to 525 bp downstream of the stop codon. We observed that Pet309-3xHA co-migrates with the mitoribosome in *cox1Δ* mitochondria, indicating that the *COX1* mRNA is not required for Pet309 association with the mitoribosome (Figure 2A).

**Figure 2.**
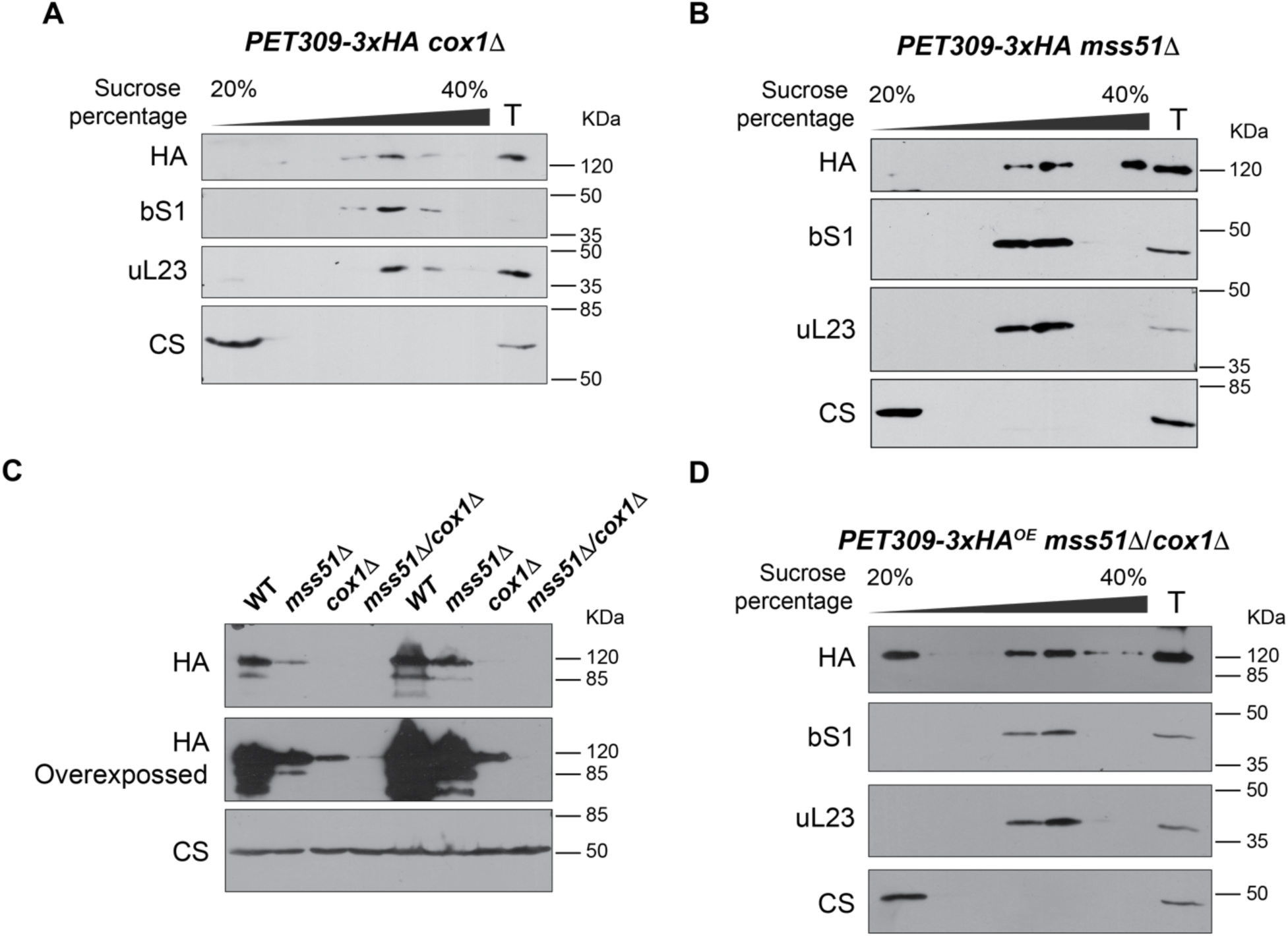
Pet309 constitutively associates to the mitoribosome. Mitochondria bearing Pet309-3xHA with the mutants A) *cox1Δ* or *mss51Δ* were lysed and separated through a sucrose gradient as in Figure 1. Fractions from the gradient were analyzed by western blot. C) Steady state levels of Pet309-3xHA in the indicated mutants were analyzed by western blot. D) Western blot analysis from a sucrose gradient separation of the mitochondrial lysate from Pet309-3xHA mitochondria bearing the double mutants *cox1Δ*, *mss51Δ*. Citrate synthase (CS) was used as loading control or a protein independent form the respiratory chain function. bS1, small subunit protein; uL23, large subunit protein; T, Total fraction, equivalent to 7% of the load. Uncropped blots in Figure S4.

In contrast to Pet309, Mss51 has not been demonstrated to interact with the *COX1* mRNA, and the mechanisms of how Mss51 activates Cox1 synthesis remains unclear (Zambrano et al., 2007, Zamudio-Ochoa et al., 2014). We hypothesize that Mss51 enable Pet309 association with the mitoribosome, explaining the requirement of Mss51 for *COX1* mRNA translation. To test this hypothesis, we expressed Pet309-3xHA into a *mss51Δ* strain, and analyzed its migration in sucrose gradients. Pet309-3xHA still co-migrated with the mitoribosome in the absence of Mss51, demonstrating that Mss51 is not required for Pet309 association with the mitoribosome (Figure 2B). Although, steady state levels of Pet309 were already low in *mss51*Δ or *cox1*Δ strains, in the double *mss51*Δ/*cox1*Δ mutant levels of Pet309 were nearly undetectable (Figure 2C). One explanation was that Pet309 loses its stability in the absence of Mss51 and the *COX1* mRNA due to its failure to associate with the mitoribosome, as observed in the *ρ°* strain (Figure 1C). Therefore, we evaluated the association of Pet309 to the mitoribosome in the double mutant *mss51Δ*/*cox1Δ*, but this time by using the *PET309-3xHA^OE^* strain to allow for Pet309 detection. Surprisingly, Pet309-3xHA^OE^ still co-migrated with the mitochondrial ribosome, indicating that neither Mss51 nor the *COX1* mRNA are necessary for Pet309 association with the mitoribosome (Figure 2D). We observed that some Pet309-3xHA^OE^ was also present in the surface fraction, likely due to the overproduction of Pet309-3xHA and saturation of the mitoribosomes.

In conclusion, our data indicate that Pet309 has a stable and constitutive interaction with the mitoribosome, independent on the presence of Mss51 and the *COX1* mRNA. However, the steady-state levels of Pet309 are compromised in the absence of either Mss51, the *COX1* mRNA or the mitoribosomes.

### The amino-terminus region of Pet309 is necessary for its interaction with the mitoribosome and with the *COX1* mRNA

As previously indicated, Pet309 is a PPR protein that belongs to the Alpha-solenoid structured tandem repeat group of proteins (Arrias et al., 2024). Analysis of the Pet309 sequence by AlphaFold predicted the presence of at least 24 PPR motifs distributed along the entire sequence (Figure 3A) (Jumper et al., 2021). Three different modules are distinguished and seem to be separated by discrete turns. The 12 central PPRs motifs, previously characterized conformed one of the three modules (presented in red in Figure 3A) (Zamudio-Ochoa et al., 2014). This region of Pet309 forms a clear superhelical structure containing well-ordered PPR motifs, and this module’s structure was predicted by AlphaFold with high score values (Figure S1A) (Jumper et al., 2021). The amino terminal region (containing 6 PPR motifs) conformed the first module (indicated in blue in Figure 3A), where the PPR repeats are also well ordered but conform a flatter structure. The last 5 to 6 PPR motifs, comprising the Pet309 C-terminus, defined the third module (indicated in yellow in figure 3A). This module seems to be less ordered as compared to the central and N-terminus modules, with longer and shorter alpha-helixes, and in some cases with breaks on some of them.

**Figure 3.**
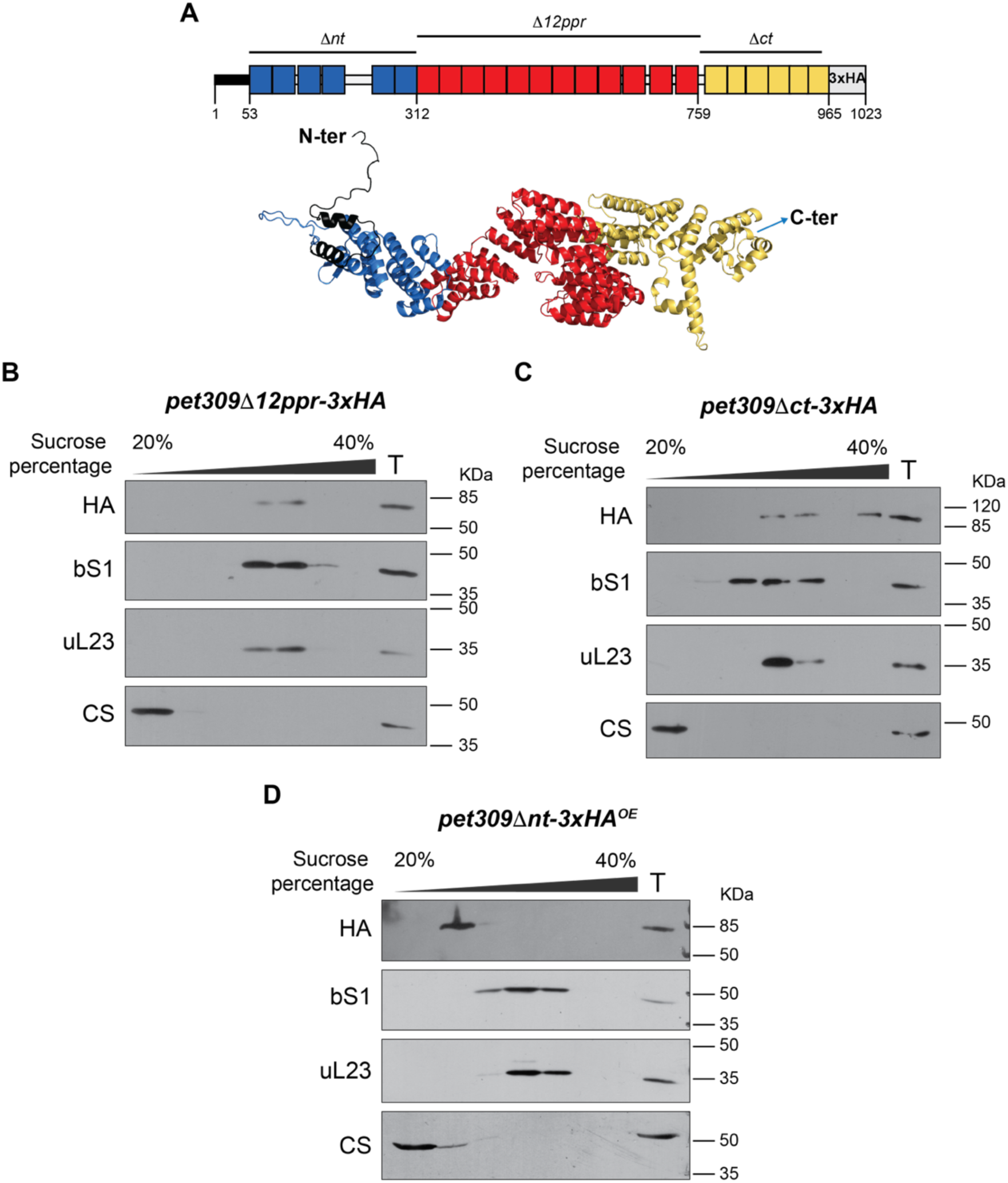
The N terminal PPR module of Pet309 mediates interaction with the mitoribosome. A) Diagram indicating the 3regions form Pet309 that were analyzed in this study. Pet309Δnt lacks residues A53 to Q311, Pet309Δ12ppr lacks residues N312 to N759 and Pet309Δct lacks residues L760 to V962. The triple hemagglutinin epitope (3xHA) was used to immunodetect Pet309 proteins with antibodies as indicated. Low panel, a model of Pet309 was downloaded from the AlphaFold database of structural modeling (Jumper, et al., 2021). The 3 proposed modules that conform Pet309 are indicated by colors: The amino terminus module, blue; the central module bearing 12 PPRs, red; the carboxy terminus module, yellow. The figure was created using Pymol. B) Western blot analysis of the sucrose gradient separation from mitochondria carrying the *pet309Δ12ppr-3xHA* B), *pet309Δnt-3xHA* C) and *pet309Δnt-3xHA* constructs. bS1, small subunit protein. uL23, large subunit protein. CS. Citrate synthase. T, Total fraction, equivalent to 7% of the load. Uncropped blots in Figure S5.

We previously demonstrated that the lack of the 12 central PPR motifs reduced Pet309 affinity for the *COX1* mRNA, abolishing translation (Zamudio-Ochoa et al., 2014). Therefore, we aimed to evaluate if Pet309 association with the mitoribosome is similarly affected by the absence of the 12 central PPR motifs. We cloned a *pet309Δ12ppr-3xHA* construct on a centromeric plasmid and transformed into *pet309Δ* cells (Zamudio-Ochoa et al., 2014). This version of the protein lacked the central module from N312 to N759. After centrifuging mitochondrial lysates on a sucrose gradient, Pet309Δ12ppr-3xHA was detected co-migrating with the mitochondrial ribosome (Figure 3B), indicating that, while the 12 central PPR motifs are essential for *COX1* mRNA translation, they do not play a role in Pet309 association with the mitoribosome.

Pet122 and Cbs2, translational activators of the *COX3* and *COB* mRNAs, respectively, interact with the mitoribosome through their C-terminal regions (Haffter et al., 1990, Krause-Buchholz et al., 2004, McMullin et al., 1990). To determine if Pet309 C-terminal domain is similarly necessary for its interaction with the mitoribosome, we generated a mutant lacking the last PPR motifs (*pet309*Δ*ct-3xHA*), just downstream of the central PPR domain, from L760 to V962. This region is part of the third PPR module of the Pet309 predicted structure (Yellow region in Figure 3A). Although the association of Pet309Δct-3xHA with the mitochondrial inner membrane was unaltered, it failed to activate *COX1* mRNA translation (Figure S1B-F). Pet309Δct-3xHA co-migrated with uL23 and bS1 in sucrose gradients, indicating that the C-terminal domain does not mediate Pet309 interaction with the mitoribosome (Figure 3C).

Since the central and C-terminal Pet309 modules were not required for ribosome association, we aimed to evaluate if the N-terminal region of Pet309 had a role on mitoribosome binding by creating a mutant without the first six PPR domains (*pet309*Δ*nt-3xHA*), from A53 to Q311. This module is predicted to have a more open conformation than the central module (blue module on Figure 3A). Like its wild-type counterpart, Pet309Δnt-3xHA remained associated with the mitochondrial inner membrane as a peripheral protein but was completely unable to activate Cox1 synthesis (Figure S2). Remarkably, Pet309Δnt-3xHA did not co-migrated with uL23 and bS1 in the sucrose gradient, and most of Pet309Δnt-3xHA was detected in surface fractions (Figure 3D). This result suggests that the ability of Pet309 to bind the mitoribosome resides within its first six predicted PPR domains.

As mentioned earlier, deletion of the 12 central PPR motifs compromised the Pet309-*COX1* mRNA binding. Therefore, we were interested in studying how deletion of the N-terminal module and the C-terminal module of Pet309 affected binding to the *COX1* mRNA. Thus, we performed and RNA-immunoprecipitation assay (Zamudio-Ochoa et al., 2014). Mitochondria bearing the *pet309Δct-3xHA* or the *pet309Δnt-3xHA* mutations were solubilized with dodecyl-maltoside. Next, the Pet309 variants were immunoprecipitated with a commercial anti-HA antibody (Figure 4A). After extensive washing, total RNA was purified from the immunoprecipitates and cDNA was obtained by reverse transcription with specific primers for *COX1* and *VAR1*. The *VAR1* gene was used as a negative control, since Pet309 does not interact with *VAR1* mRNA. The cDNA was further analyzed by PCR to amplify the *COX1* 5’ UTR or *VAR1* 5’ UTR regions. The *COX1* 5’ UTR was successfully amplified in IP fractions from both Pet309-3xHA and Pet309Δct-3xHA. As expected, the *COX1* 5’ UTR was not detected in the untagged Pet309 IP control. Additionally, *VAR1*, was not amplified in any IP fractions (Figure 4B). In contrast, interaction of Pet309Δnt-3xHA^OE^ with the *COX1* mRNA 5’ UTR was undetectable, suggesting that the amino terminal module of Pet309 is necessary for *COX1* mRNA binding (Figure 4 C, D).

**Figure 4.**
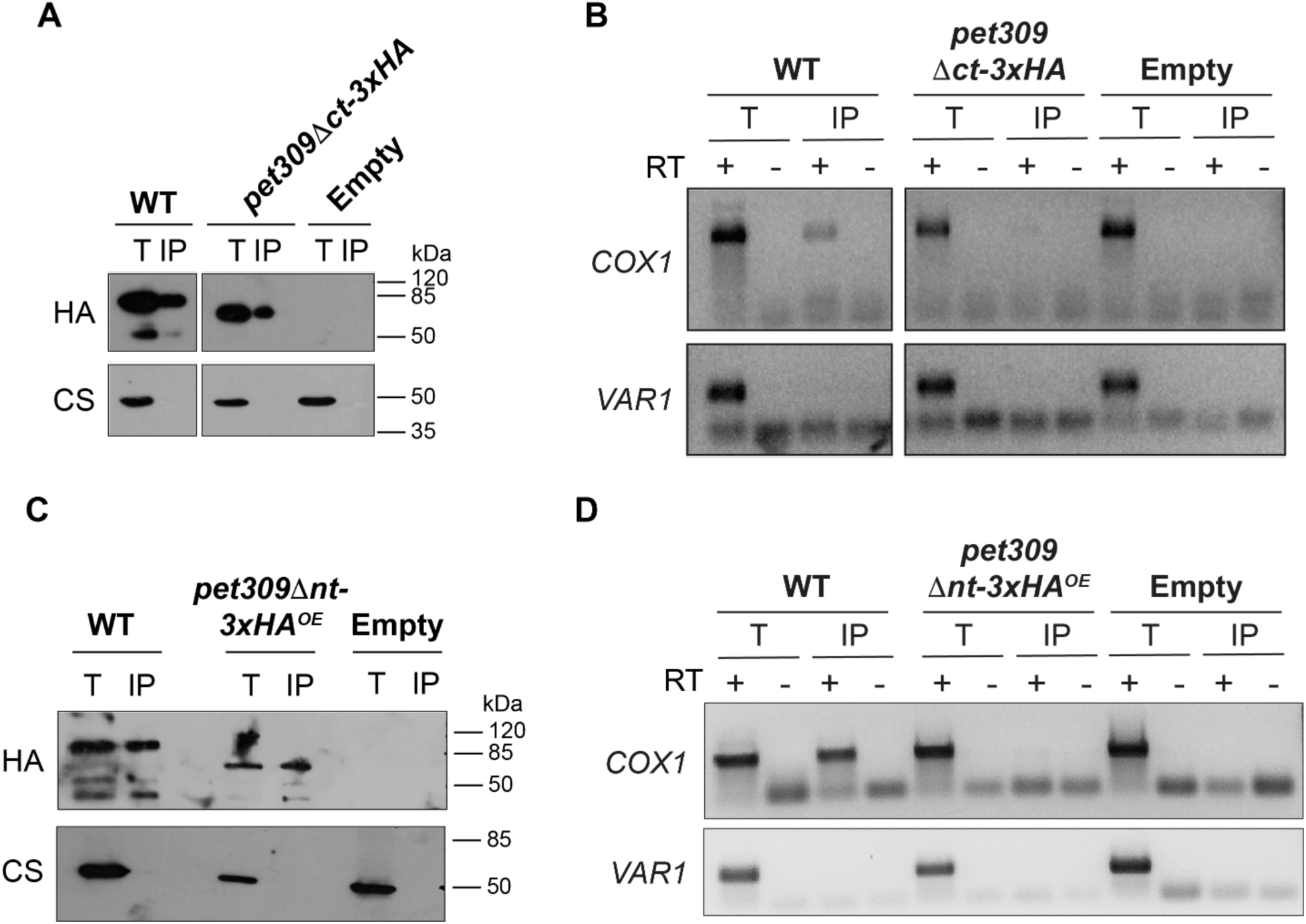
The N-terminal module of Pet309 is necessary for interaction with the *COX1* mRNA. Mitochondria from *PET309-3xHA*, A) *pet309Δct-3xHA* or C) *pet309Δnt-3xHA^OE^* and the untagged strain were lysed with digitonin. The lysates were subjected to immunoprecipitation with anti-HA antibodies coupled to agarose beads. 5% of the lysate, 25% of the immunoprecipitated (IP) and 25% of the supernatant fractions (S) were separated by SDS-PAGE and analyzed by Western blot using an anti-HA antibody (HA) to show the efficiency of immunoprecipitation. An anti-citrate synthase antibody (CS) was used as a negative control. Next, RNA was extracted from the total (T), immunoprecipitated (IP) and supernatant (S) fractions from B) *pet309Δct-3xHA* or D) *pet309Δnt-3xHA^OE^* samples. After DNase treatment, cDNA was prepared using reverse transcriptase (RT) (+) and adding primers for *COX1* and *VAR1* genes (Supplemental Table 2). Samples without RT (-) were included as a control for DNA contamination. Resulted cDNAs were used as a template for PCR amplification of *COX1* and *VAR1* genes and run in an agarose gel. Uncropped blots in Figure S5.

It is well-established that Pet309 stabilizes the *COX1* mRNA, and its overexpression leads to an accumulation of *COX1* mRNA, likely due to its capacity to protect the RNA from endogenous nucleases (Figure S3) (Zamudio-Ochoa et al., 2014). To determine if the overexpression of Pet309Δct-3xHA or Pet309Δnt-3xHA leads to *COX1* mRNA accumulation, we transformed a *pet309*Δ strain with constructs expressing *pet309*Δ*ct-3xHA* (or *pet309*Δ*nt-3xHA*), *PET309-3xHA* (as a positive control), or an empty vector (as a negative control), from high- or low-copy expression vectors. We purified total RNA from all strains and the accumulation of the *COX1* mRNA was analyzed by northern blot using ^32^P-labeled oligonucleotides specific for *COX1*, *COX2* (as negative control), and *15S* rRNA (as loading control). Overexpression of Pet309-3xHA and Pet309Δct-3xHA led to an accumulation of *COX1* mRNA, whereas *COX2* and *15S* rRNA levels remained unchanged, demonstrating a specific effect of Pet309 overexpression on the *COX1* mRNA accumulation (Figure 5A, B). These data indicated that deletion of the last four PPR motifs in Pet309 does not abolish its interaction with the *COX1* mRNA. In contrast, over expression of *pet309*Δ*nt-3xHA* failed to accumulate *COX1* mRNA (Figure 5C, D). The inability of Pet309Δnt-3xHA to bind and stabilize the *COX1* mRNA is not due to alterations in its mitochondrial localization and association with the mitochondrial inner membrane, which is similar to the wild type Pet309-3xHA (Figure S2).

**Figure 5.**
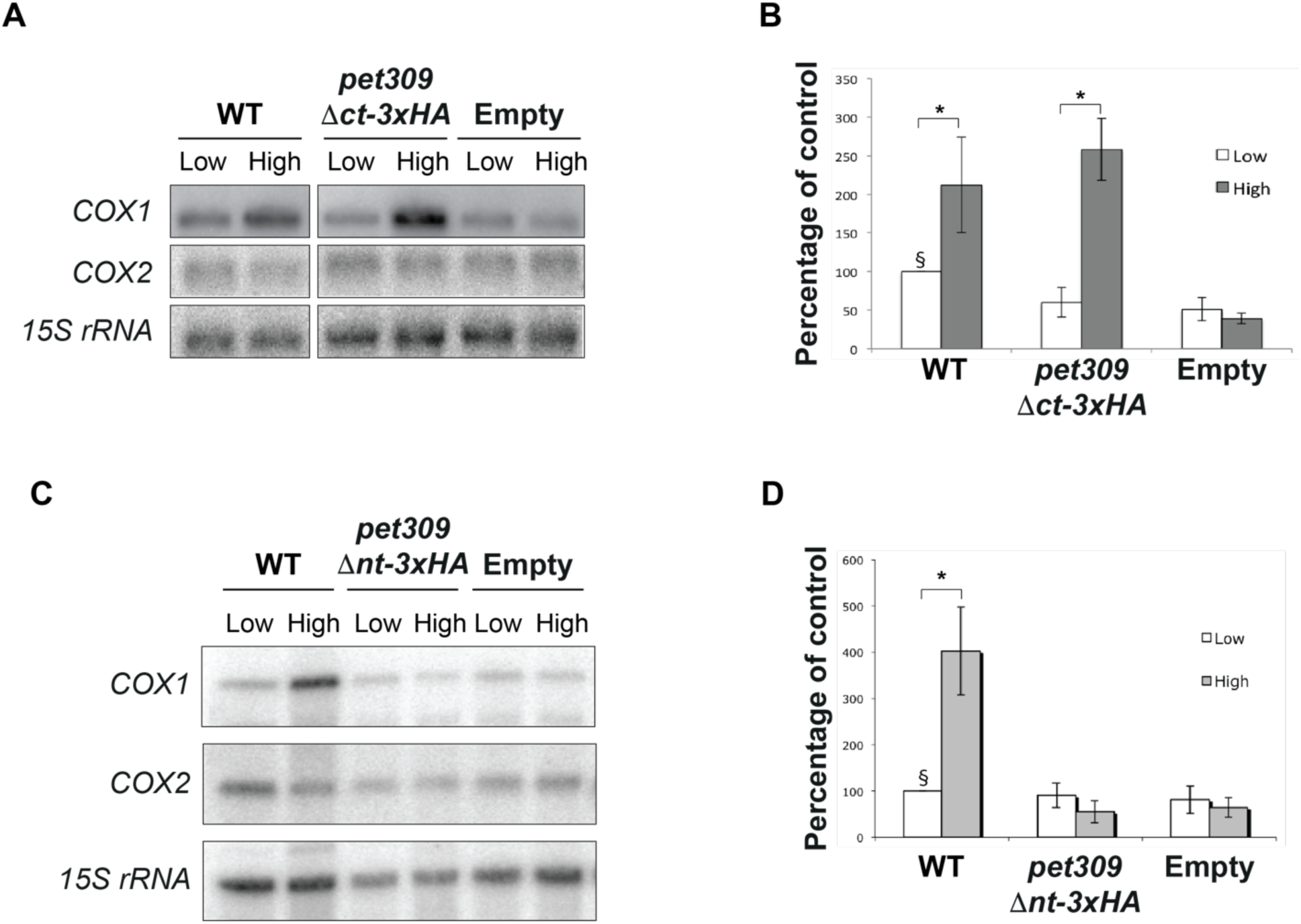
The N terminal module from Pet309 is involved in stabilization of the *COX1* mRNA. The *PET309-3xHA*, *pet309Δct-3xHA* A) *pet309Δnt-3xHA* B) were cloned on centromeric plasmid for low copy number (Low) and 2μ plasmid for high copy number (High) and transformed into a *pet309Δ* strain. An empty plasmid was also transformed and used as negative control. 10μg of total RNA was analyzed by Northern blot. As a negative control, total RNA from *pet309Δ* strains bearing empty plasmids was also analyzed. The membrane was hybridized with ^32^P labeled probes complementary to *COX1*, *COX2* and *15S rRNA* probes. Quantification of the *COX1* signals from three independent experiments are represented in a bar graph. *COX1* signals were normalized against *15S* rRNA and the result from *PET309-3xHA* in low copy plasmid was taken as 100% (§). (*) Statistical analysis was made by using a 2-way ANOVA with Bonferroni post-hoc test (*p* < 0.05).

Taken together, these data indicate that the first six PPR motifs (N terminal module) of Pet309 are critical for binding the *COX1* mRNA and to associate with the mitoribosome, while the central and C-terminal end modules are not required for these processes. However, the three modules are indispensable for translational activity of Pet309.

### Mss51 interacts with the mitoribosome independently of the presence of Pet309 or the *COX1* mRNA

Mss51 coordinates Cox1 synthesis and assembly by acting on the *COX1* 5’ UTR and physically interacting with the Cox1 peptide (Barrientos et al., 2004, Perez-Martinez et al., 2003, Perez-Martinez et al., 2009). However, the mechanisms by which Mss51 activates *COX1* mRNA translation are yet to be understood. It was previously reported that a population of Mss51 comigrated in sucrose gradient centrifugations with the translational machinery (De Silva et al., 2017, Fontanesi et al., 2010, Mays et al., 2019, Verma et al., 2021). However, these comigrations were mainly observed in C*c*O assembly mutants (De Silva et al., 2017, Fontanesi et al., 2010, Mays et al., 2019, Verma et al., 2021). Moreover, Mss51 was undetectable in the MIOREX complexes (Kehrein et al., 2015). Therefore, Mss51 association with the mitoribosome under wild-type conditions remains unclear. To illuminate Mss51’s role as a translational activator, we focused on evaluating whether Mss51 associates with the mitoribosome in a wild type context. We used a strain expressing Mss51-3xHA endogenously in its original locus which was previously shown to not disturb its respiratory growth (Perez-Martinez et al., 2003). As expected, most of the Mss51-3xHA population was found in surface fractions of the sucrose gradient, but a small amount was detected co-migrating with the mitoribosome (Figure 6A). In contrast, in a *ρ°* strain lacking mitochondrial DNA (and thereof mitoribosomes) Mss51 was solely detected in surface fractions, indicating that a small fraction of Mss51 is associated with mitoribosomes (Figure 6B). Since only a small population of Mss51 co-migrated with the mitoribosome, maybe the Mss51-mitoribosome interaction could be transient and occur only when the *COX1* mRNA is being translated. To determine if Mss51 association with the ribosome depends on active translation of the *COX1* mRNA, we analyzed Mss51-3xHA migration in the *cox1*Δ strain used earlier (Figure 2A). Interestingly, Mss51-3xHA co-migrated in sucrose gradients with bS1 and uL32, indicating that Mss51 interacted with the mitoribosome independently of the presence of the *COX1* mRNA (Figure 6C). Previous reports demonstrated that Pet309 and Mss51 physically interact (Zamudio-Ochoa et al., 2014), and since we found that Pet309 constitutively interacts with the mitoribosome, we hypothesized that the Mss51-mitoribosome interaction could be mediated by Pet309. To address this hypothesis, we analyzed the association of Mss51 with the mitoribosome in a mutant lacking Pet309. Mss51-3xHA maintained its co-migration with bS1 and uL23 in the absence of Pet309 (Figure 6D), and the same result was obtained after simultaneous deletion of Pet309 and the *COX1* gene (Figure 6E). Taken together, these results indicate that a small fraction of Mss51 constitutively interacts with the mitoribosome, independently on the presence of the *COX1* mRNA and Pet309.

**Figure 6.**
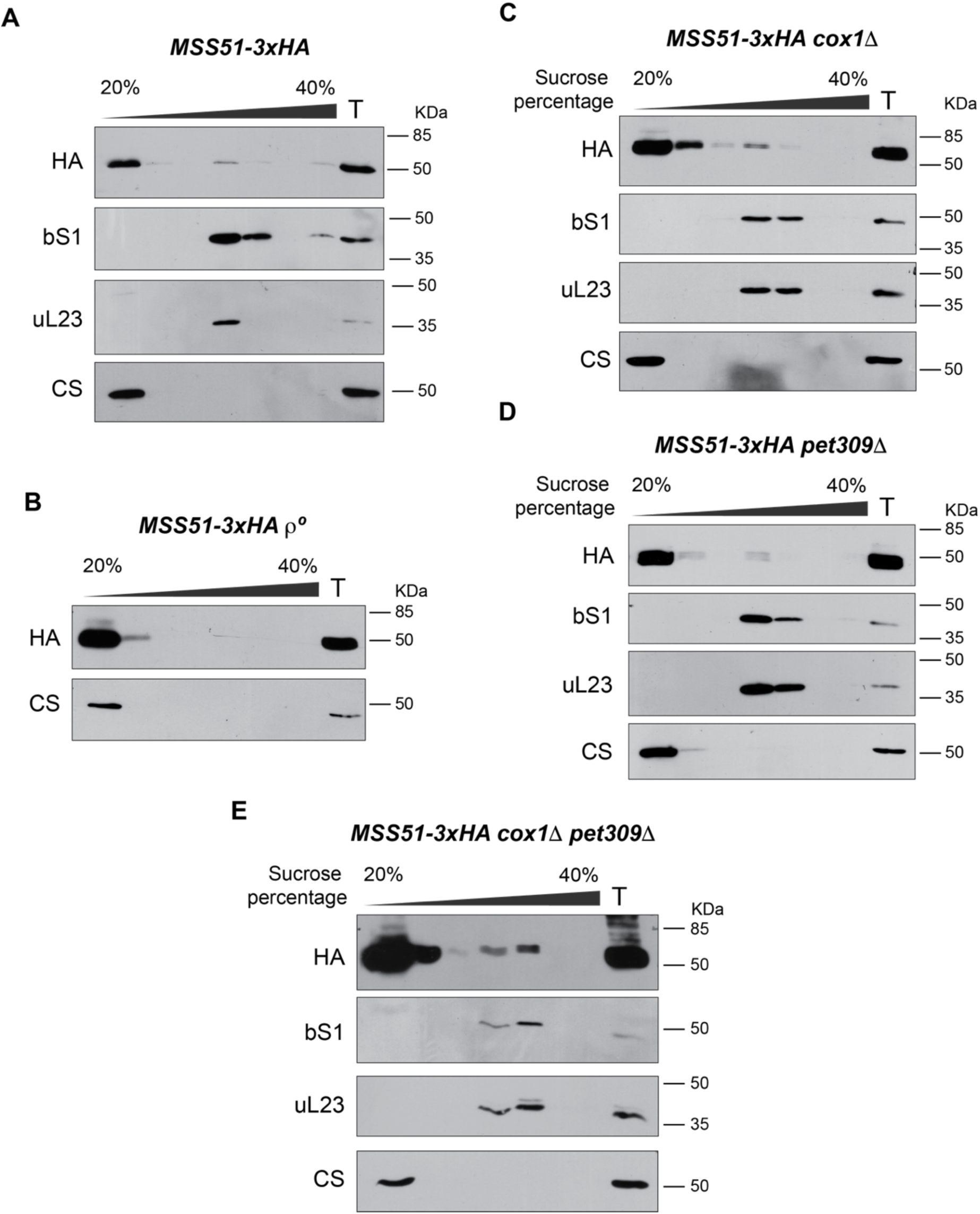
Mss51 constitutively interacts with the mitoribosome. Mitochondria from *MSS51-3xHA* cells carrying A) WT mitochondrial DNA or B) devoid of mtDNA (*ρ°*) were solubilized and separated by ultracentrifugation on a sucrose gradient. The resultant seven fractions from this gradient were analyzed by Western blot with the indicated antibodies. Similar experiments were carried out with mitochondria bearing C) a deletion of the *COX1* gene (*cox1Δ*), D) the mutant *PET309* gene (*pet309Δ*), or E) the double mutant *cox1Δ pet309Δ*. Uncropped blots in Figure S6.

## DISCUSSION

### Pet309 as a constitutive interactor of the mitoribosome

Pet309 and Mss51 were first described 30 years ago (Decoster et al., 1990, Manthey and McEwen, 1995), yet their mechanisms of translational activation remain unclear. In this study, we demonstrated that Pet309 and Mss51 interact with the mitoribosome independently of each other and of the *COX1* mRNA. While Pet309 was not detected as a ribosome-free protein, only a minor proportion of Mss51 interacted with the mitoribosome. This aligns with previous findings, as Mss51 also interacts with Cox1 assembly intermediates (Fontanesi et al., 2011, Mick et al., 2010, Perez-Martinez et al., 2003, Pierrel et al., 2007).

The instability of Pet309 in the absence of mitoribosomes (in the *ρ°* strain) suggests a constitutive association of Pet309 with the translational machinery. This decrease in steady-state levels of Pet309 is unlikely due to changes in its synthesis, as cytosolic translation is unaffected in *ρ°* strains (Couvillion et al., 2016). Mitoribosomal proteins are prone to degradation when the ribosome is unassembled (Graack and Wittmann-Liebold, 1998), a phenomenon we also observed for bS1 and uL23 in our *ρ°* strains. Our findings suggest that Pet309 is more than a transiently associated ribosome-accessory protein; it may resemble a mitoribosomal subunit. This is supported by the observation that Pet309’s association with the mitoribosome in MIOREX complexes is resistant to high ionic strength, while the helicase Mss116 dissociates under similar conditions (Kehrein et al., 2015).

Intriguingly, Pet309 can interact with the mitoribosome even in the absence of Mss51 or the *COX1* mRNA, although its stability is compromised if either is missing. In fact, the combined absence of both Mss51 and *COX1* mRNA further destabilizes Pet309. This may suggest that a complete *COX1* mRNA translation machinery is necessary to protect Pet309 from degradation.

Pet309’s association with the mitoribosome occurs independently of the *COX1* mRNA, implying that certain mitoribosomes may be preloaded with Pet309 to initiate *COX1* mRNA translation. This raises the question of whether Pet309 is a marker of specialized ribosomes dedicated to Cox1 synthesis or mitochondria-encoded C*c*O subunits. The concept of heterogeneous ribosome populations preferentially translating specific mRNAs is gaining acceptance (for a review, see (Guo, 2018)). Indeed, the yeast mitoribosome cryo-EM structure reveals a non-identified protuberance above the mRNA exit canyon, which might represent different translational activators that label a subset of mitoribosomes (Desai et al., 2017).

Pet309 is composed almost entirely of PPR motifs. We previously demonstrated that eliminating a single PPR motif in the central module rendered a non-functional protein, although it could still bind *COX1* mRNA (Tavares-Carreon et al., 2008). Although Pet309’s target sequence has not been identified, two sites – at −356 to −335 and −76 to −55 nt from the *COX1* mRNA AUG start codon – are predicted as binding sites (De Silva et al., 2017). Deletion of a single PPR or multiple repeats, as in this study, likely disrupts the RNA-protein recognition code. AlphaFold predicts approximately 24 PPR motifs in Pet309, forming three distinct modules. The central module, composed of 12 PPR motifs, is the most structured, forming a compact superhelical structure. The N-terminal module, containing 6 PPR motifs, is predicted to have a more open, planar structure, while the C-terminal end module consists of 3 well-structured PPR motifs followed by 3 disorganized PPR motifs, with varying alpha-helix lengths and turns within some helices (Figure 3A and Figure S1A). We demonstrated that the central and C-terminal end modules of Pet309 are dispensable for its binding to *COX1* mRNA and the mitoribosome, whereas the N-terminal module is essential for both. Although Pet309Δnt-HA lacks association with the mitoribosome, it was detected in the second fraction of the gradient, suggesting its presence in a complex with other proteins, possibly protecting it from degradation a 900 kDa complex containing Pet309 has been described previously (Krause et al., 2004), which includes the *COB* mRNA translational activator Cbp1 (Islas-Osuna et al., 2002, Mittelmeier and Dieckmann, 1990).

Deleting Pet309’s N-terminal module also abolished its interaction with the *COX1* mRNA. Pet309’s ribosomal location may be essential for *COX1* mRNA interaction, supported by findings that the *COX1* mRNA associates with the mitoribosome even in Pet309’s absence (De Silva et al., 2017), and that mitochondrial mRNAs are channeled from transcription to translation by specialized factors (Kehrein et al., 2015). Nam1, which associates with mitoribosomes and interacts with RNA Polymerase and various translational activators, including Pet309, may be one such channeling factor (Bryan et al., 2002, Kehrein et al., 2015, Markov et al., 2009, Naithani et al., 2003, Rodeheffer et al., 2001). We believe the lack of interaction between Pet309Δnt and the *COX1* mRNA is due to the mutant’s altered location, distant from translation machinery and channeling factors.

Beyond the initial six PPR motifs, other PPR mutants analyzed in this, and prior studies did not affect Pet309’s ability to bind the *COX1* mRNA and the mitoribosome, though they did impair Cox1 synthesis (Zamudio-Ochoa et al., 2014). Two potential explanations exist: first, that missing PPR motifs disrupt Pet309’s ability to accurately recognize the *COX1* sequence, mispositioning the mitoribosome; or second, that these deletions affect Pet309’s interaction with other proteins. The DEAD-box helicase Mss116 interacts with Pet309 and is proposed to unwind *COX1* secondary structures to facilitate sequence-specific binding by Pet309. Interestingly, Mss116 mutants affecting helicase activity did not disrupt its interaction with Pet309, although *COX1* translation was impaired (De Silva et al., 2017).

### Pet309 and Mss51 role in activating *COX1* mRNA translation

Mss51 is another essential translational activator of *COX1* mRNA, but unlike Pet309, it plays a dynamic role in C*c*O biogenesis by coordinating Cox1 synthesis and assembly (Barrientos et al., 2004, Mick et al., 2010, Perez-Martinez et al., 2009). In this study, we found that Mss51 interacts with the mitoribosome, although this interaction appears weak and/or transient, as only a minor fraction of Mss51 was detected in ribosomal fractions and it was absent in MIOREX complexes (Kehrein et al., 2015). Interestingly, Mss51 remains associated with the mitoribosome even in the absence of the *COX1* mRNA, suggesting that, a subset of Mss51 is constitutively associated with the translational machinery independently of *COX1* mRNA translation. This indicates that Mss51, like Pet309, may interact with the mitoribosome prior to Cox1 synthesis.

Both, Pet309 and Mss51 activate *COX1* mRNA translation, but their mitoribosome interactions are independent.

Based on these results and prior studies on Cox1 biogenesis, we propose the following model for the *COX1* mRNA translation (Figure 7):

1. *<u>Preloading Phase</u>. Pet309, Mss51 and Mss116 associate with the mitoribosome before the COX1 mRNA is loaded.* Our data suggest that all of Pet309 is bound to the mitoribosome, while Mss51 exists in two populations: one engaged in Cox1 assembly and a smaller fraction associated with the mitoribosome. Pet309 behaves like a ribosomal subunit, as it withstands high ionic strength conditions, via its N-terminal module, whereas Mss116 acts as a ribosomal accessory, detaching under similar conditions (Kehrein et al., 2015).
2. *<u>mRNA Binding and Positioning</u>*: *The helicase Mss116 unwinds the secondary structure of the approaching COX1 mRNA to enable Pet309 binding. Mss51 induces a conformational change on Pet309 enabling it to recognize its specific sequence on the 5’ UTR and to position the mitoribosome active site on the AUG start codon.* Two putative sequences on the *COX1* 5’UTR could be targets of Pet309 (De Silva et al., 2017). The helicase Mss116 assist the binding process, probably by unwinding secondary structures in the mRNA to facilitate Pet309 to localize its cognate sequence (De Silva et al., 2017). However, these interactions alone may be insufficient for accurate Pet309’s sequence binding and correct ribosome positioning, as seen in *mss51*-null mutants that exhibit abnormal *COX1* mRNA translation dependent on Pet309. (Zambrano et al., 2007). Mss51 could facilitate this process by altering Pet309’s conformation, modifying the interaction of Pet309 with the *COX1* 5’ UTR. This aligns with previous findings showing that Pet309’s immunoprecipitation is inefficient without Mss51, likely due to conformational changes affecting epitope accessibility, though Pet309 still interacts with the *COX1* mRNA (Zamudio-Ochoa et al., 2014). Indeed, PPR proteins frequently undergo conformational shifts (Ban et al., 2013, Ke et al., 2013, Shen et al., 2016, Yin et al., 2013).
3. *<u>Translation elongation</u>. Pet309 remains bound to the mitoribosome and to the COX1 mRNA throughout translation but does not actively participate in elongation, while Mss51 and Mss116 are still required*. Evidence shows that Pet309 binds to the *COX1* mRNA even when elongation is inhibited by puromycin (Zamudio-Ochoa et al., 2014). While Mss51 and the helicase Mss116 target additional *COX1* coding sequence regions, Pet309 acts exclusively on the 5’ UTR (De Silva et al., 2017, Perez-Martinez et al., 2003, Perez-Martinez et al., 2009, Zamudio-Ochoa et al., 2014). In this context, we propose that during elongation Pet309’s interaction with *COX1* mRNA may be nonspecific since it lacks coding sequence recognition.
4. *<u>Post-Translation and Cox1 Assembly</u>. After COX1 mRNA translation, Pet309 remains attached to the ribosome, while Mss51 engages in the assembly of the nascent Cox1 peptide, where it may change its own conformation and/or association with the ribosome* (Barrientos et al., 2004, Mick et al., 2010, Perez-Martinez et al., 2003). At least three different Mss51 conformations have been proposed (Fontanesi et al., 2011, Mick et al., 2011, Mick et al., 2010), though it is unclear which conformation remains ribosome-bound or if additional conformations exist. After assisting with Cox1 assembly, Mss51 adopts a translationally inactive conformation until new rounds of translation initiation take place (step 1).

**Figure 7.**
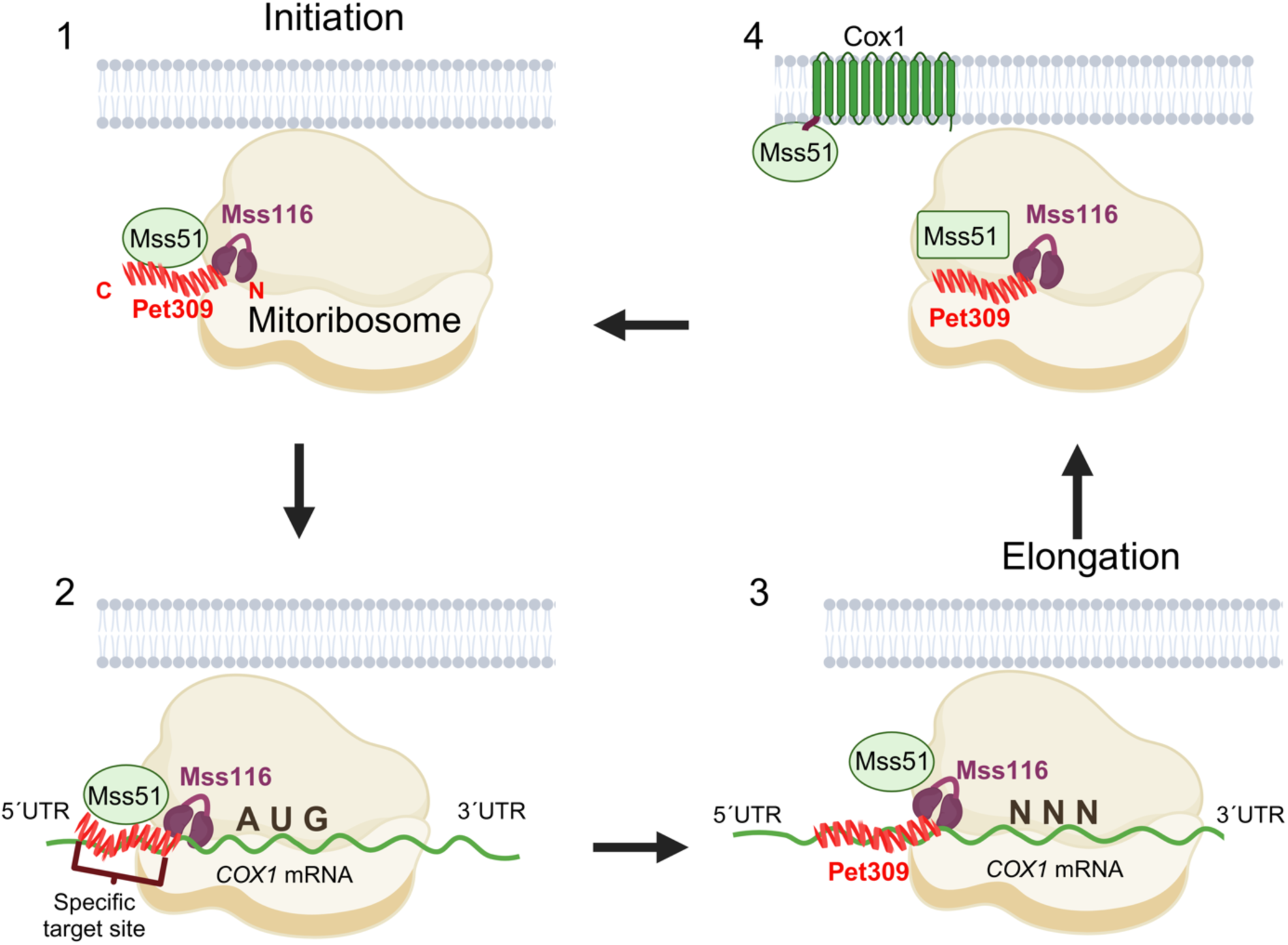
Model for the mechanisms of Pet309 and Mss51 on translation of the *COX1* mRNA. See text in Discussion section for details.

Interestingly, Mss51’s role in respiratory regulation is also present in mammals. Mammalian Mss51 is expressed mainly in skeletal muscle where it functions as a respiratory repressor. Mss51 knock-out mice show increased respiration, insulin sensitivity and obesity resistance (Rovira Gonzalez et al., 2019). This poses Mss51 as a potential therapeutic target for obesity and type-2 diabetes prevention (Ali et al., 2024). Both yeast and mammalian Mss51 contain conserved zinc-finger domains (Baleva et al., 2021), though the exact function, whether DNA binding or protein interaction, remains unexplored. Further research is needed to clarify Mss51’s mechanisms in C*c*O biogenesis in yeast and respiratory regulation in mammals.

Many questions remain to be answered in future studies, such as: How does Pet309 specifically recognize the *COX1* mRNA among other mitochondrial mRNAs? How do Pet309, Mss51 and Mss116 associate to the ribosome during translation termination and ribosome recycling? What exact role does Mss51 play in aiding Pet309 and the ribosome to locate the *AUG* start codon? Are there similarities between yeast Mss51 and its mammalian counterpart’s mechanisms?

## Materials and Methods

### Yeast strains, media, and genetic methods

The *S*. *cerevisiae* strains used in this study are listed in Supplemental Table S1. Standard genetic methods and media recipes were as previously described (Burke D, 2000). Complete fermentable media were YPD or YPRaf (containing 2% glucose or 2% raffinose). Minimal media contained 0.67% yeast nitrogen base, 2% glucose or 3% ethanol/3% glycerol, and Complete Supplement Mixtures (CSMs) purchased from Bio 101 (Vista, CA) and ForMedium (UK). Gene deletion constructs with *KANMX4*, *LEU2*, or *URA3* cassettes were generated by PCR.

### Mitochondrial transformation, and integration of the cox1Δ construct into ρ^+^ mtDNA

A plasmid containing the *COX1* gene, including 1365 bp upstream of the 5’ UTR and 990 bp of the 3’ UTR was digested with Pac1 and religated to create a deletion of the complete *COX1* ORF, together with 787 bp of the 5’ UTR and 525 bp of the 3’ UTR, to create plasmid pXPM75. A fragment of the *COX2* gene was amplified by PCR and cloned into the *ClaI-HindIII* sites in pXPM75 to create plasmid pCB14. This plasmid was transformed on Nab69 *ρ°* by high-velocity microprojectile bombardment (Camacho-Villasana et al., 2024). Mitochondrial transformants were identified by their ability to rescue respiratory growth when mated to a *ρ^+^* strain bearing the A114 to stop mutation in the mitochondrial *cox2* gene. The *cox1Δ* construct was integrated by homologous recombination into *ρ^+^* mtDNA by isolating cytoductants issued from crosses of the transformant to strain TF258 (Perez-Martinez et al., 2009).

### Cloning of the pet309Δnt-3xHA and pet309Δct-3xHA constructs

The plasmids used and generated during this work are derivatives of pXP96, pXP97 and pXP104 (Tavares-Carreon et al., 2008), and contain the *PET309-3xHA* sequence, including 310 and 205 nt of the *PET309* 5′ and 3′ UTRs. Pet309 mutants were constructed similarly as previously reported *(Zamudio-Ochoa et al., 2014)*. *PET309Δnt* lacks residues A53 to Q311, and *PET309Δct* lacks residues L760 to V962. Both mutants were generated by fusion PCR (Ho et al., 1989). Site-directed mutagenesis by overlap extension using the polymerase chain reaction, using Accuzyme DNA polymerase (Bioline) and pXP97 as the DNA template. *PstI/EcoRI*-digested PCR products were cloned into similarly digested pXP96. The *Xba*I -*Xho*I DNA fragments were ligated into pXP97 to generate the ARS/CEN, low-copy-number plasmids. For high-copy-number plasmids, the *XbaI- ClaI* fragments from mutant-bearing pXP96 plasmids were ligated into the pXP104 plasmid.

### Sucrose fractionation of mitochondrial lysates

This protocol was previously reported on (Garcia-Guerrero et al., 2018). A sample of 500 μg of mitochondrial protein was incubated with lysis buffer (1% digitonin, 10 mM MgOAc, 50 mM NaCl, 20 mM HEPES-KOH, pH 7.4, 1 mM PMSF) for 30 min on ice. The mitochondrial lysate was clarified by centrifugation at 16,200 *x g* for 10 min. The supernatant of the centrifugation was then loaded into a discontinuous gradient of 40, 30, and 20% sucrose containing 0.1% digitonin, 10 mM MgOAc_2_, 20 mM DTT, 10 mM Tris-HCl, pH 7.4, and 0.5 mM PMSF. The sucrose gradients were ultracentrifuged at 145,000 g in a SW-55Ti rotor for 2 hr at 4 °C. Fractions of 600 ml were TCA-treated for protein precipitation. Proteins were resolved by SDS-PAGE, transferred to PVDF membranes, and detected by immunoblotting with the indicated antibodies.

### RNA immunoprecipitation assay

This protocol was done as previously reported (Zamudio-Ochoa et al., 2014). Briefly, 1 mg of mitochondrial protein was lysed with 500 μl of 0.7% n-Dodecyl β-D-maltoside, 100 mM NaCl, 20 mM Tris-HCl pH 7.4, 200 U of RNaseOUT (Invitrogen), and protease inhibitors (Roche). The cleared lysate was incubated with an anti-HA high-affinity antibody (Roche) coupled to protein A-sepharose (GE Healthcare). The immunoprecipitate was washed and one-fourth of the supernatant and precipitate fractions were saved for western blot analysis. The remainder was used for RNA extraction with Trizol reagent. 20 ng of RNA were treated with 1 unit of DNase I (Invitrogen) for 15 min at 25 °C. The cDNA was prepared by addition of primers for *COX1*, *COX3*, *VAR1 ATP8* or *ATP6* in the presence of SuperScript III Reverse Transcriptase (Invitrogen). This cDNA was used as template for PCR reactions to amplify the *COX1* or *COX3* 5′ UTRs. For primer list, Supplemental Table S2.

### Northern blot

Total RNA was extracted from yeast cells using the RNeasy Mini kit (Qiagen). 10 μg of total RNA was separated by denaturing agarose gel electrophoresis and transferred to Hybond XL membrane (GE Healthcare). Membranes were probed with radioactively labeled probes recognizing *COX1* exon 4, *COX2*, and *15S* rRNA. Blots were analyzed with a Typhoon 8600 PhosphorImager (GE Healthcare) and quantitated with ImageQuaNT.

### Membrane protein analyses

100 μg of mitochondria was resuspended in 400 μl of 0.1 M Na_2_CO_3_, vortexed and incubated on ice for 60 minutes. The sample was centrifuged at 100,000 *x g* for 10 minutes, and the supernatant was precipitated with 10% trichloroacetic acid.

For separation of mitochondria in membrane and soluble fractions, 100 μg of mitochondria were resuspended in 100 mM NaCl 100 mM, Hepes HEPES 20mM and sonicated. The samples were centrifuged at 100,000 *x g* for 10 minutes, and the supernatant was precipitated with 10% trichloroacetic acid.

### Proteinase K digestion of mitochondria

Samples of 100 μg of mitochondria were suspended in HEPES-KOH 20 mM pH 7.4 with or without sorbitol 0.6 M. The samples were incubated on ice with 100 μg/ml of proteinase K. Digestion was terminated with 2 mM phenylmethylsulfonyl fluoride.

### Image digitalization and processing

Original films were scanned, and edited brightness and contrast by Adobe Photoshop software following all scientific ethical criteria. Original scans are shown in Figures S4-7.

### Proofreading

The ChatGPT application was used for proofreading, followed by careful review from all authors.

## Acknowledgements

We thank Thomas D. Fox for the gift of antisera; Thomas D. Fox, Gabriel del Río-Guerra and Teresa Lara-Ortiz for the gift of yeast strains. We thank Laura Ongay-Larios, Guadalupe Codiz-Huerta and Minerva Mora-Cabrera, from UBM-IFC for plasmid sequencing. We thank Ulrik Pedroza-Dávila for careful reading of the manuscript. This work was supported by the research grant from Programa de Apoyo a Proyectos de Investigación e Innovación Tecnológica, (PAPIIT), UNAM [IN2223623 to XP-M] and Consejo Nacional de Ciencia y Tecnología (CONACyT) [CB284514 to XP-M]. Biorender agreement number for Figure 7: DL27HWG37M.

## Author contributions

XP-M conceived and coordinated the study, and together with AG-G and AZ-O wrote the manuscript. In addition, performed the experiment from Supplementary S3. AZ-O performed the experiments from figures 1-3, 5, 6, S1 and S2, and constructed yeast mutants. AG-G analyzed the results and performed Figure S2. YC-V constructed yeast strains and plasmids, performed the experiments from Figure 4 and prepared Figure 7. All authors analyzed the results and approved the final version of the manuscript.

## Data and resource availability

All relevant data and resource can be found within the article and it’s supplementary information.

## Supplemental Material

**Supplemental Table 1.** *S. cerevisiae* Strains Used in This Study.

| Strain | Genotype | Reference |
| --- | --- | --- |
| AGG54 | *Mata, ura3-52, ade2, arg8::hisG, lys2, leu2-3, 112, mss51Δ::LEU2, PET309::3xHA nuc1Δ::KanMX4 [ρ<sup>+</sup>]<sup>a</sup>* | (Zamudio-Ochoa et al., 2014) |
| FTC25 | *Mataα, ura3Δ, ade2 PET309-3xHA nuc1Δ::KanMX4 [ρ<sup>+</sup>]<sup>a</sup>* | (Zamudio-Ochoa et al., 2014) |
| FTC26 | *Mataα, ura3-52, leu2-3, 112 lys2 arg8::hisG pet309Δ::LEU2 nuc1Δ::KanMX4 ρ<sup>+</sup>, [ΔΣai]<sup>b</sup>* | (Zamudio-Ochoa et al., 2014) |
| YC133 | *Mataα, ura3Δ, ade2 nuc1Δ::KanMX4 [ρ<sup>+</sup>]<sup>a</sup>* | (Zamudio-Ochoa et al., 2014) |
| SB7 | *Mataα, ura3Δ, ade2 MSS51-3xHA, [ρ<sup>+</sup>]<sup>a</sup>* | (Perez-Martinez et al., 2003) |
| SB5 | *Mataα, ura3Δ, ade2 PET309-3xHA, [ρ<sup>+</sup>]<sup>a</sup>* | (Zamudio-Ochoa et al., 2014) |
| XPM232 | *Mataα, ura3-52, leu2-3, 112 lys2 arg8::hisG pet309Δ::LEU2 [ρ<sup>+</sup>, ΔΣai]<sup>b</sup>* | (Zamudio-Ochoa et al., 2014) |
| JPM16 | *Mataα, ura3Δ, ade2, pet309Δ::URA3, MSS51-3xHA [ρ<sup>+</sup>]<sup>a</sup>* | This work |
| JPM22 | *Mataα, ura3Δ, ade2, PET309-3xHA, mss51Δ::KanMX4 [ρ<sup>+</sup>]<sup>a</sup>* | This work |
| AZO53 | *Mataα, ura3-52, leu2-3, 112 lys2 arg8::hisG pet309Δ::LEU2, [ρ<sup>+</sup>, cox1Δ::ARG8<sup>m</sup>]<sup>a</sup>* | This work |
| AZO86 | *Mataα, ura3Δ, ade2 MSS51-3xHA, [ρ<sup>+</sup>, cox1Δ::ARG8<sup>m</sup>]<sup>a</sup>* | This work |
| YC140 | *Mataα, ura3Δ, ade2, pet309::URA3, MSS51-3xHA, [[ρ<sup>+</sup>, cox1Δ::ARG8<sup>m</sup>]<sup>a</sup>* | This work |
| YC135 | *Mataα, ura3-52, leu2-3 112, lys2 arg8::hisG, pet309Δ::LEU2, mss51::KANMX4 [ρ<sup>+</sup>, cox1Δ::ARG8<sup>m</sup>]<sup>a</sup>* | This work |

<sup>a</sup> Mitochondrial genome is shown in brackets
<sup>b</sup> Mitochondrial *COX1* gene has no introns
All strains are isogenic or congeneric to D273-10b

**Supplemental Table 2.**
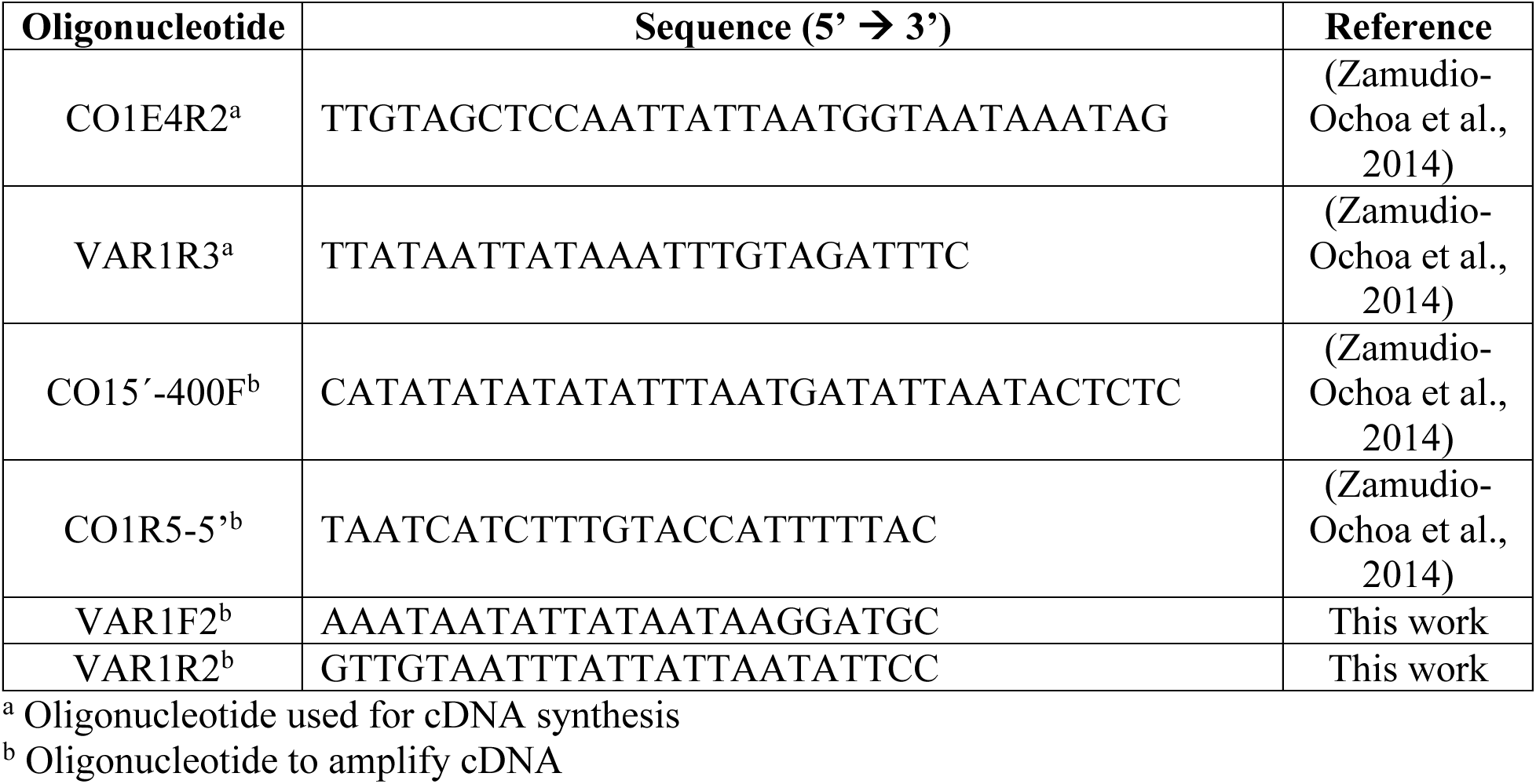
Oligonucleotides Used in RT-PCR Experiments.

**Figure S1.**
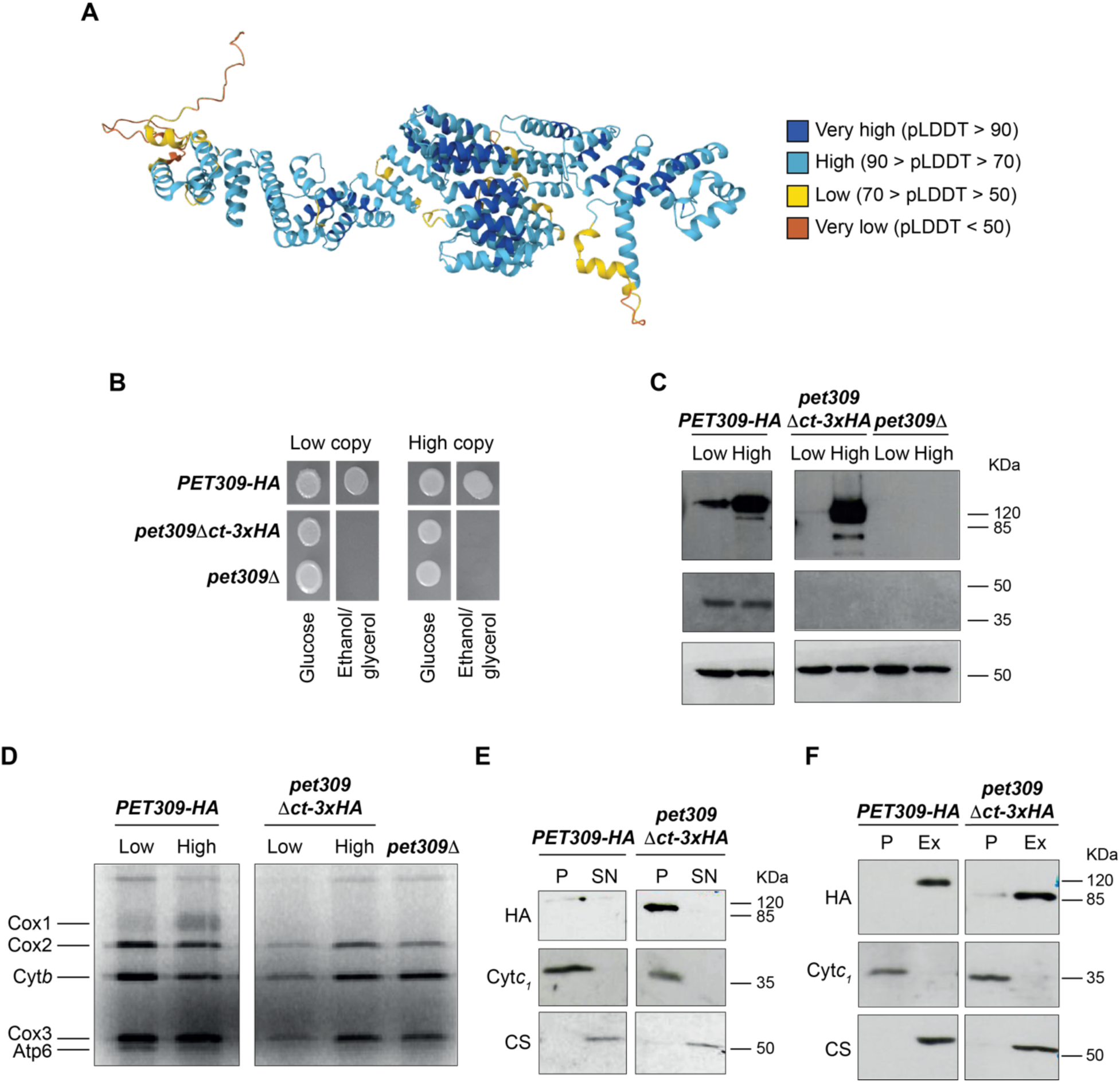
A mutant lacking the C-terminus module of Pet309 localized to mitochondria like a wild type Pet309 protein. A) The Pet309 full sequence was analyzed in the AlphaFold software. The model is shown with the color code used by the program to indicate values of confidence. The model obtained a pTM value of 0.73 (Jumper et al., 2021). B) The *pet309Δct-3xHA* and the *PET309-3xHA* constructs were cloned in either centromeric (Low) or 2μ (High) yeast expression plasmids and were transformed in cells lacking the endogenous *PET309* gene (pet309Δ was transformed with empty plasmid, as negative control). Growth on fermentative or respiratory media was tested for 3 days at 30°C. C) The same strains were used for western blot analysis of mitochondria using the indicated antibodies. D) Cells were incubated with ^35^S-methionine and cycloheximide to inhibit cytoplasmic ribosomes. Mitochondrial translation products were analyzed by SDS-PAGE and autoradiography. E) Mitochondria from *PET309-3xHA* or *pet309Δct-3xHA* strain were sonicated to break the membranes. The membrane (P) and the soluble fractions (S) were separated by ultracentrifugation, resolved by SDS-PAGE and analyzed by Western blot. Anti-Cyt*c_1_* antibody was used as a control of a membrane protein and anti-CS (citrate synthase) antibody was used as a control of a soluble protein. F) Mitochondria from *PET09-3xHA* and *pet309Δct-3xHA* strain were treated with Na_2_CO_3_ to extract non-integral proteins from the mitochondrial membranes. After ultracentrifugation, membranes (pellet, P) and extracted and soluble proteins (EX) were resolved with SDS-PAGE and analyzed by Western blot. The Anti-Cyt*c_1_* antibody was used as a control of an integral membrane protein and the anti-CS antibody as a control of a soluble protein. Uncropped blots in Figure S7.

**Figure S2.**
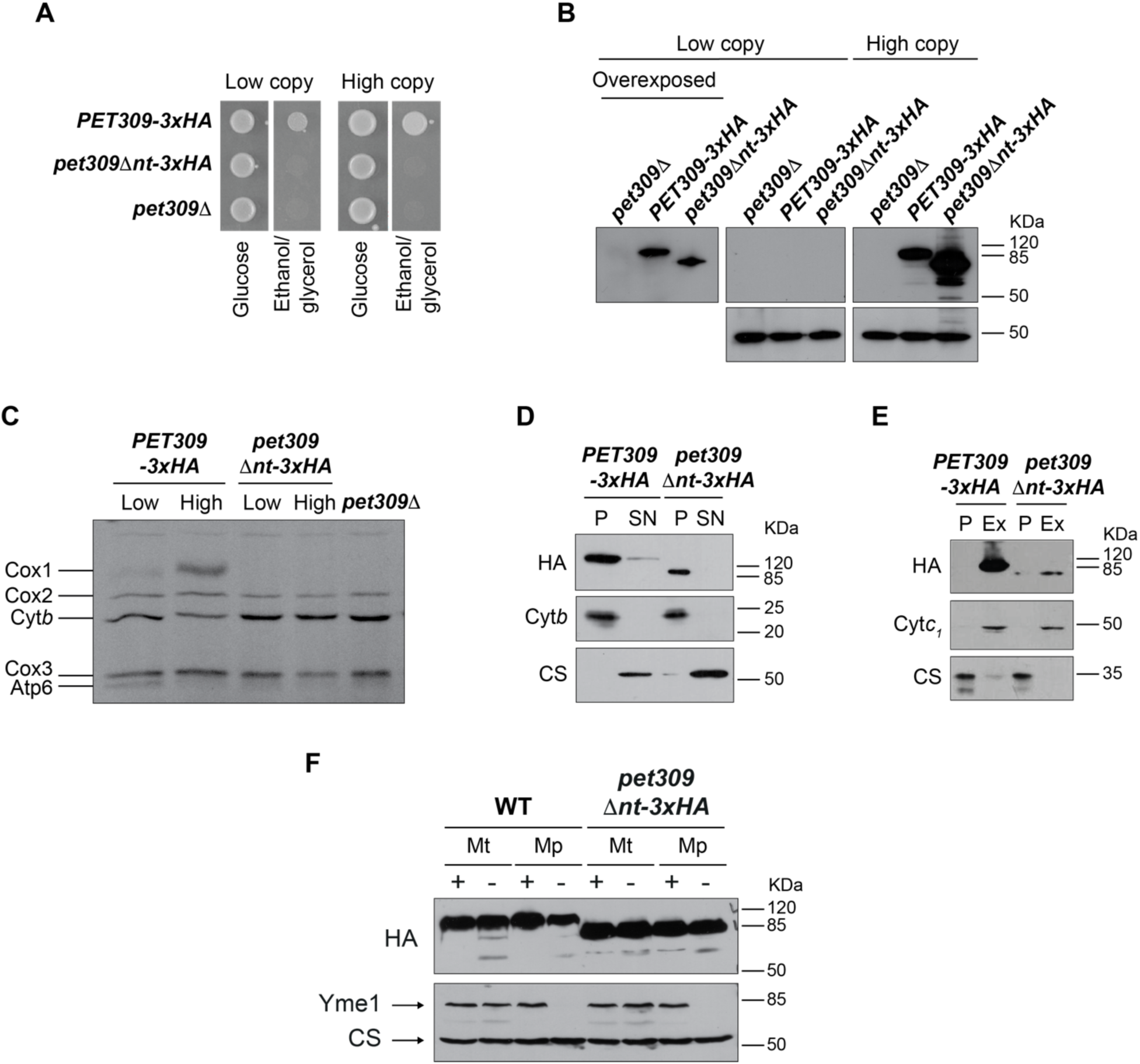
A mutant lacking the N-terminus module of Pet309 localized to mitochondria like a wild type Pet309 protein. A) The *pet309Δnt-3xHA* and the *PET309-3xHA* constructs were cloned in either centromeric (Low) or 2μ (High) yeast expression plasmids and transformed in cells lacking the endogenous *PET309* gene. *pet309Δ* strain was transformed with empty plasmid. Growth on fermentative or respiratory media of the indicated cells was tested for 3 days at 30°C. B) The same strains were used for western blot analysis of mitochondria. C) Cells were incubated with ^35^S-methionine and cycloheximide as in Supplementary Figure S1. D) Mitochondria from *PET309-3xHA* or *pet309Δnt-3xHA* strain were sonicated to break the membranes. The membrane (P) and the soluble fractions (S) were separated by ultracentrifugation, and analyzed by Western blot as in Supplementary Figure S1. D) Mitochondria from Pet309-3xHA and *pet309Δnt-3xHA* strain were treated with Na_2_CO_3_ to extract non-integral proteins from the mitochondrial membranes. After ultracentrifugation, membranes (pellet, P) and extracted and soluble proteins (EX) were resolved with SDS-PAGE and analyzed by Western blot. E) Mitochondria from the *PET309-3xHA* or *pet309Δnt-3xHA* strains were incubated in hypotonic (Mitoplasts, Mp) or isotonic (Mitochondria, Mt) buffer in the absence (-) or presence (+) of Proteinase K. Samples were resolved by SDS-PAGE and analyzed by Western blot using anti-HA, anti-Yme1 and anti-Citrate synthase (CS) antibodies. Uncropped blots in Figure S7.

**Figure S3.**
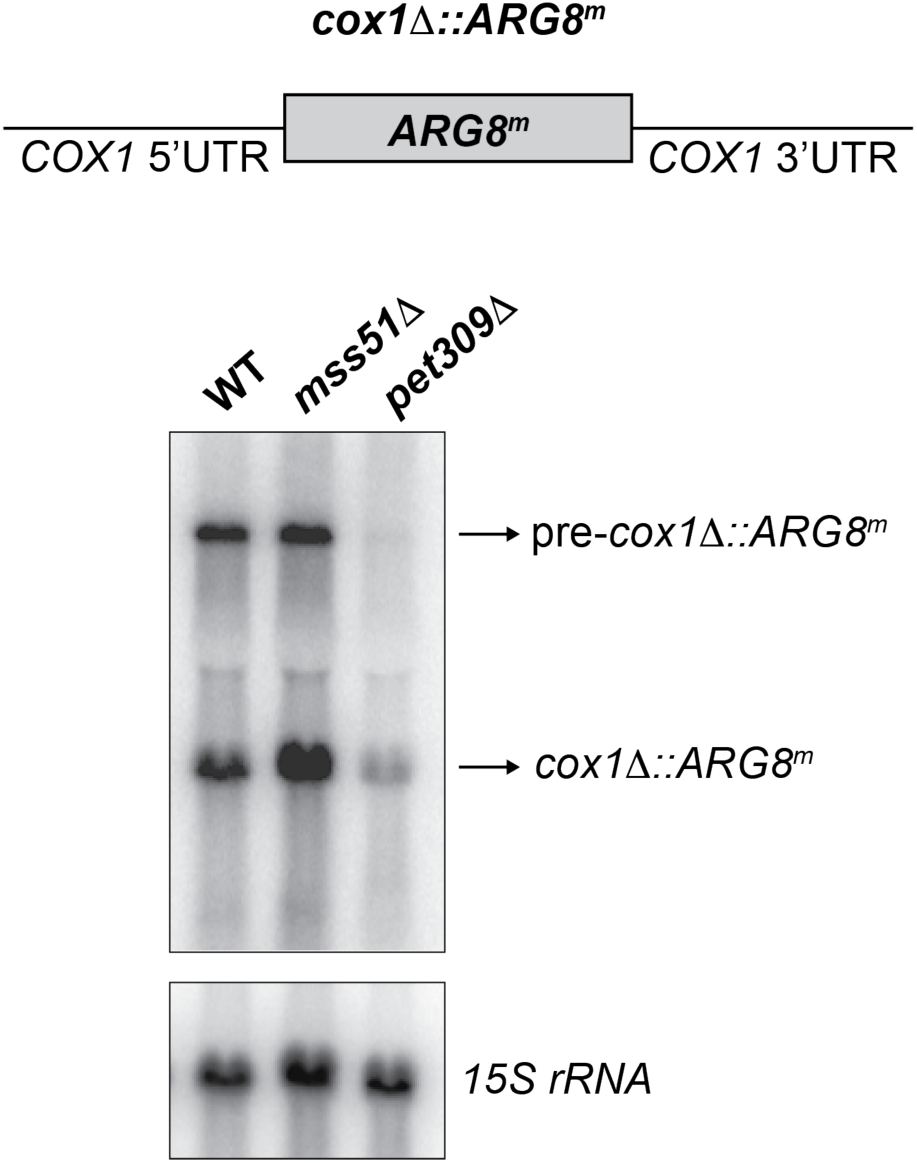
Pet309, and not Mss51, is necessary for stability of the *cox1Δ::ARG8^m^* mitochondrial mRNA. 10μg of total RNA from WT, *mss51Δ* or *pet309Δ* cells were isolated and separated by agarose electrophoresis. ^32^P-radiolabeled probes for *ARG8^m^* and the *15S* rRNA were used.

**Figure S4.**
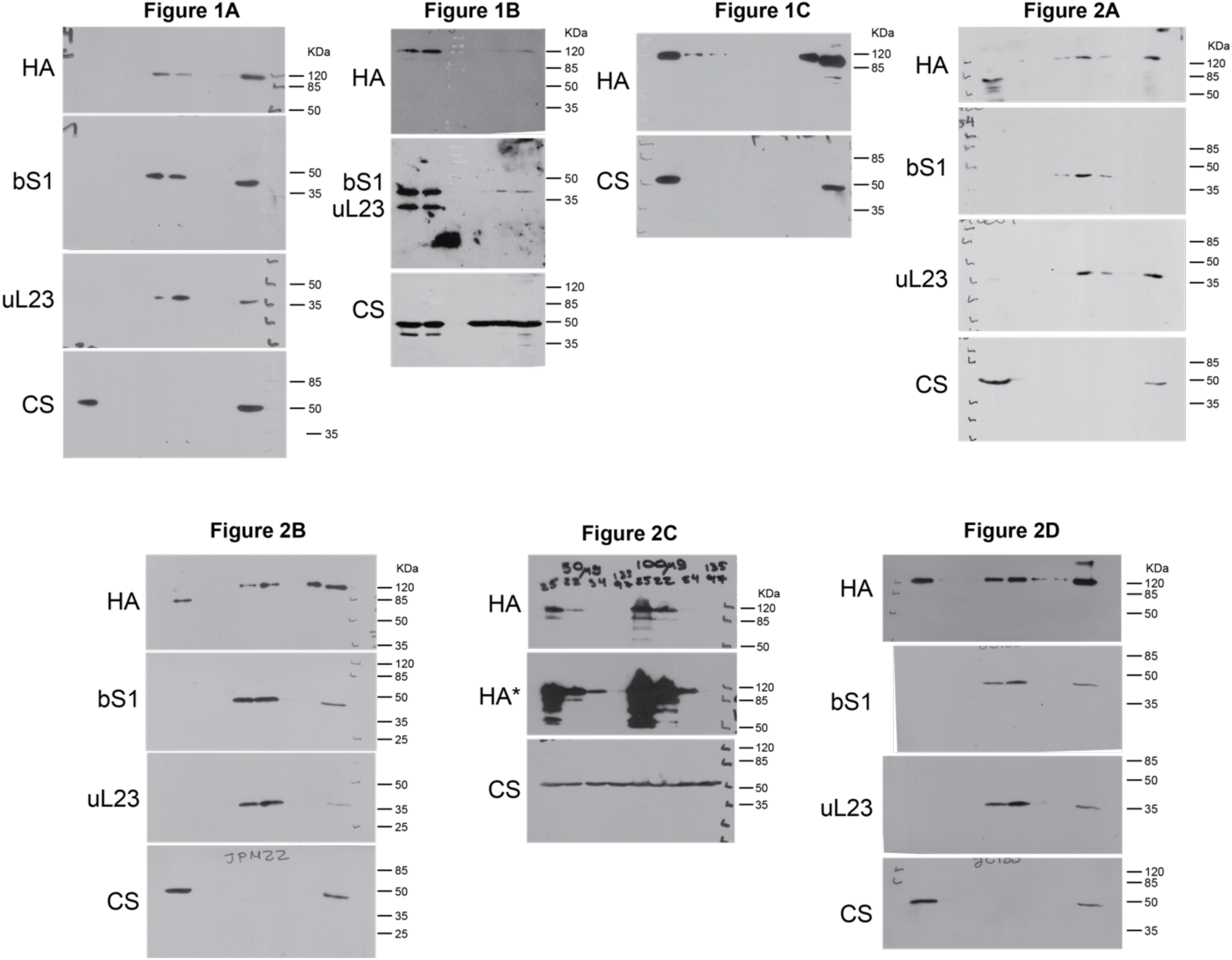
Uncropped blots from Figures 1 and 2. Blots were imaged by film exposure and scanned for digitalization. Used antibodies indicated on each blot.

**Figure S5.**
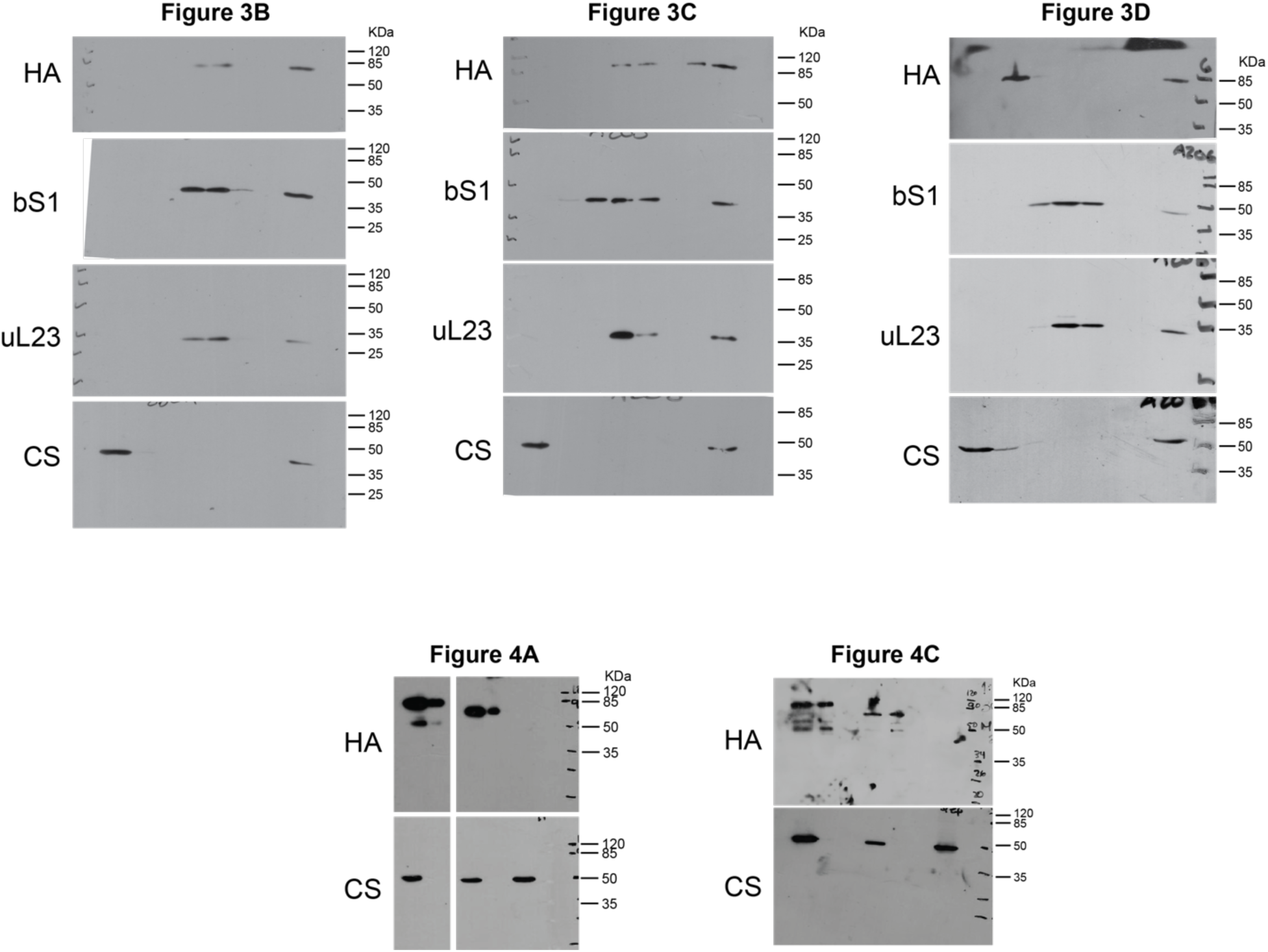
Uncropped blots from Figures 3 and 4. Blots were imaged by film exposure and scanned for digitalization. Used antibodies indicated on each blot.

**Figure S6.**
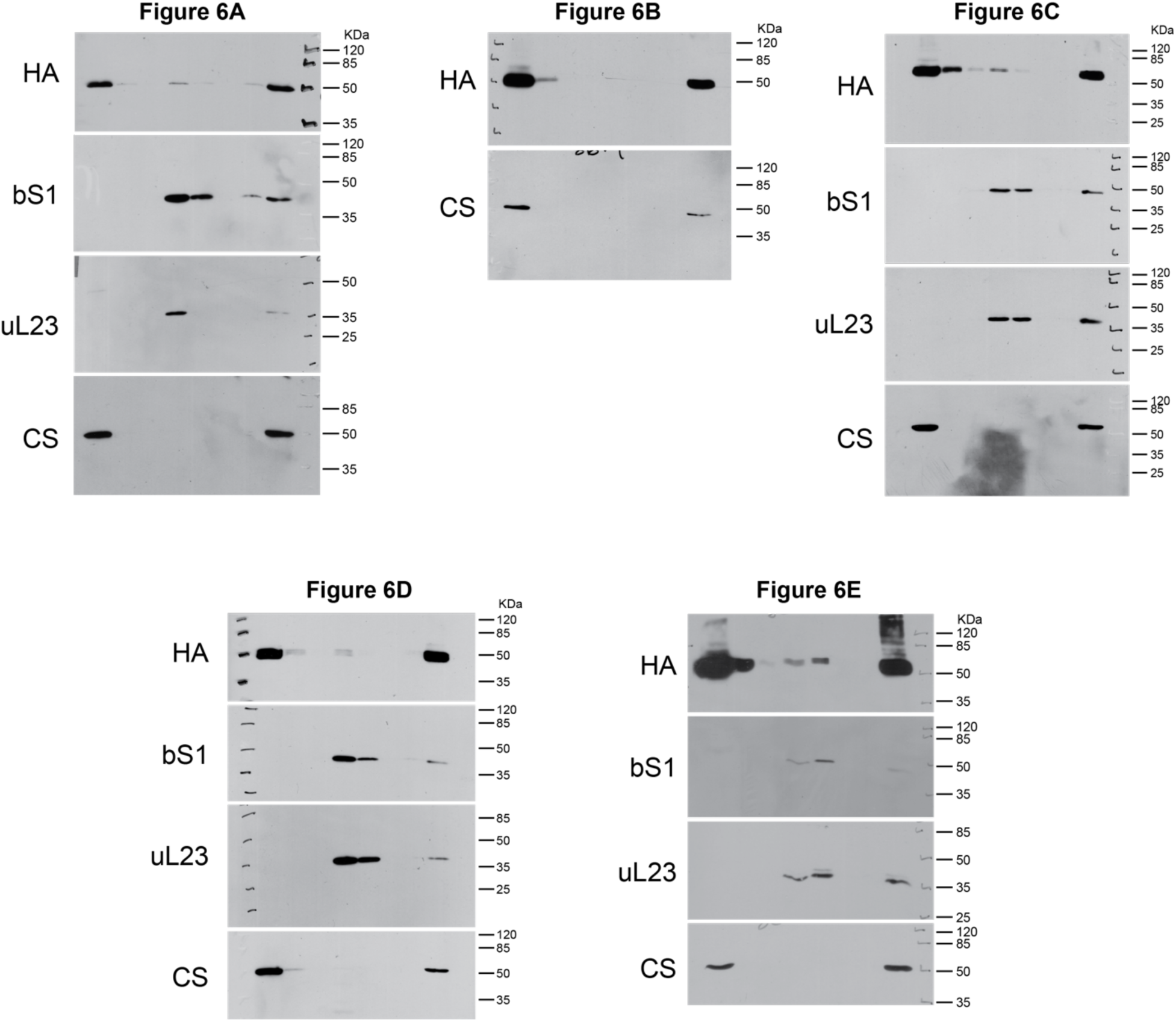
Uncropped blots from Figure 6. Blots were imaged by film exposure and scanned for digitalization. Used antibodies indicated on each blot.

**Figure S7.**
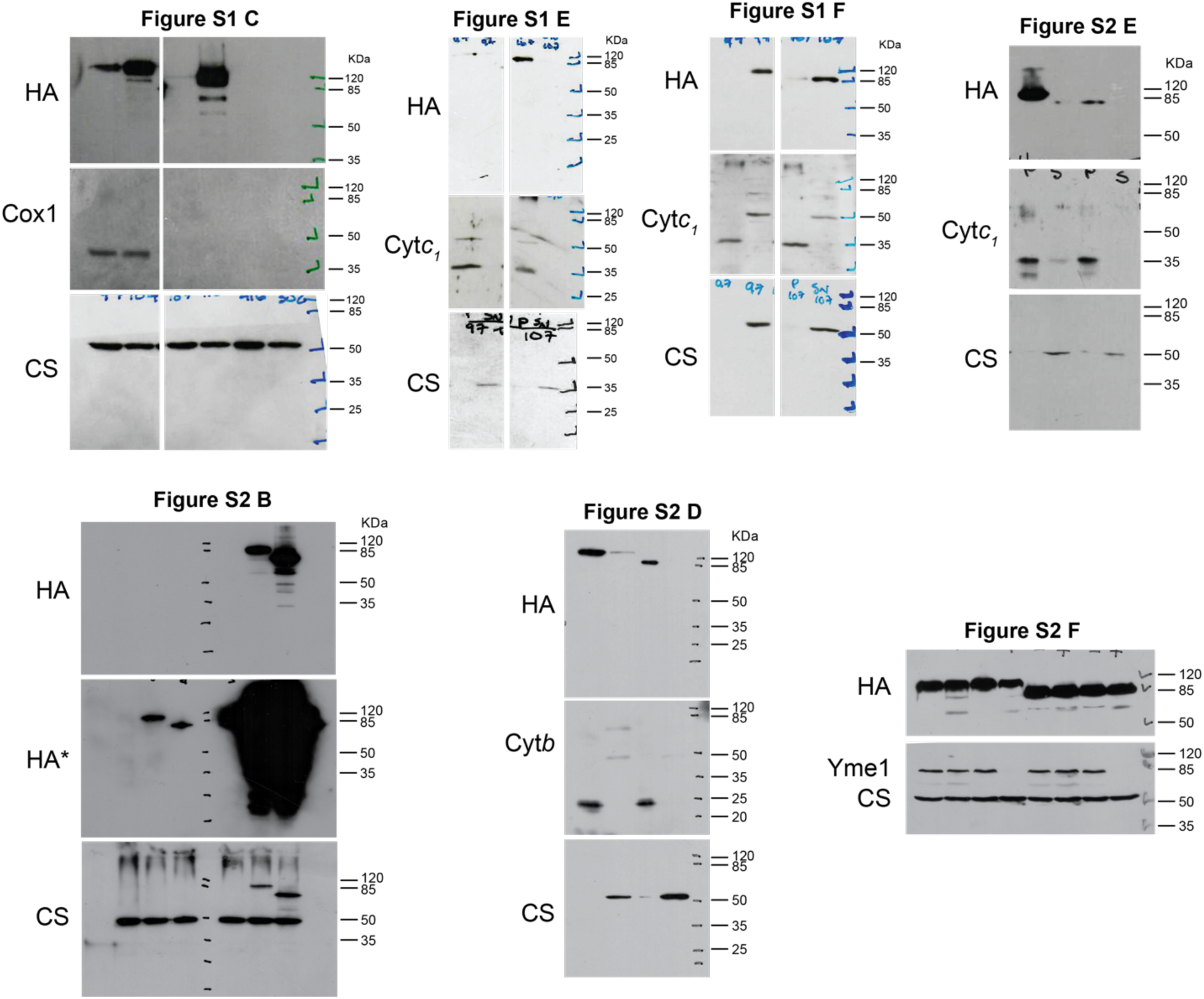
Uncropped blots from Figures S1 and S2. Blots were imaged by film exposure and scanned for digitalization. Used antibodies indicated on each blot.

